# Phase resetting of in-phase synchronized Hodgkin-Huxley dynamics under voltage perturbation reveals reduced null space

**DOI:** 10.64898/2026.03.21.713085

**Authors:** Rahul Gupta, Karmeshu, R. K. Brojen Singh

## Abstract

Voltage perturbations to a repetitively firing Hodgkin-Huxley (HH) model of neuronal spiking in the bistable regime with coexisting limit cycle and stable steady node can either lead to the spike’s phase resetting or collapse to the stable steady state. The latter describes a non-firing hyperpolarized quiescent state of the neuron despite the presence of constant external current. Using asymptotic phase response curve (PRC), the impact of voltage perturbations on a repetitively firing HH model is studied here while it is diffusively coupled to another HH model under identical external stimulation. It is observed that the pre-perturbation state of synchronization and the coupling strength critically determine the PRC response of the perturbed HH dynamics. Higher coupling strengths of perfectly in-phase (anti-phase) synchronized HH models shrink (expand) the combinatorial space of perturbation strengths and the oscillation phases causing collapse to the quiescent state. This indicates reduced (enlarged) basin of attraction, viz. the null space, associated with the steady state in the HH phase space. The findings bear important implications to the spiking dynamics of diverse interneurons, as well as special cases of pyramidal neurons, coupled through electrical synapses via. gap junctions, and suggest the role of gap junction plasticity in tuning vulnerability to quiescent state in the presence of biological noise and spikelets.

## Introduction

The biological neurons are perfect examples of natural oscillators. Given a sufficiently-strong constant external stimulating current, a typical neuron repeatedly fires action potential (AP). The Hodgkin-Huxley (HH) model [1, 2] is a widely-recognized biophysically-detailed model of AP generation [3–6] in the soma and axon initial segments (AIS) of various neurons [7–9]. Bistable behaviour of the HH model has been extensively studied using mathematical and computational approaches [10–17]. Within a certain suprathreshold range of external current, a stable focus co-exists with the stable limit cycle in the phase space as a result of subcritical Hopf and saddle-node bifurcations. The stable limit cycle represents repeated AP formation, whereas stable focus represents a hyperpolarized quiescent state. A neuron at the stable focus cannot fire APs despite the presence of strong external current. While understanding that HH model of neuronal spiking is best suited to describe neuron’s somatic and AIS compartments, for the sake of simplicity, we refer to the HH model as the HH neuron under point approximation in the following text.

Experimental studies in a variety of neurons have confirmed the presence of bistable phenomenon in their spiking behaviour. In rat cerebellar Purkinje cells, blockade of the hyperpolarization-activated current (*I*_*h*_) reveals bistability between tonic action potential firing and a quiescent hyperpolarized state [18]. Brief current pulses can cause switch to the quiescent state. Similarly, in vivo recordings show Purkinje cells [19] and spiny striatal cells [20] transitioning between up (spiking) and down (silent, hyperpolarized) membrane potential states in response to synaptic input or current pulses. Similar effects of *I*_*h*_ current controlling the state transitions have been observed in spinal motoneurons [21]. Optogenetic/inhibitory manipulations and electrical stimulation have been shown to produce extended silencing or prolonged shifts into hyperpolarized states in cortex and other regions [22]. Modelling study in leech heart interneuron [23] has also shown existence of hyperpolarized quiescent states under bistability.

In the context of natural oscillators, Winfree proposed a methodology to study the phase resetting behavior of a system with a limit cycle attractor [24–27]. Perturbations of different magnitudes are applied to the system at different phases of the oscillation. The new phases of the oscillation restoration are recorded after some integer multiples of the time period. In this way, an asymptotic phase resetting curve (PRC) is obtained between the old and the new phases. Often small-magnitude perturbations lead to only minute changes in the new phase. However, a dramatic change in the new phase for a sufficiently large perturbation suggests crossing of the system across a null space in the dynamical phase space. According to the Winfree’s limit cycle theory [25, 28], the null space is the subset of the dynamical phase space not contained in the attractor basin manifold of the limit cycle. Nudging the system into the null space leads to annihilation of the rhythmic oscillation. Thus, the new phase is undefined or indeterminate in the PRC. Therefore, phase resetting behaviour of a HH-type neuron repeatedly firing APs becomes intriguing in the bistable regime.

Using Winfree’s methodology, Best computationally investigated the asymptotic PRC of the HH neuron in the presence of a constant external current under the bistable regime [29]. Best chose to perturb membrane potential, as this could also be performed in an experimental setup with squid axon. Using phase-stimulus plot, he mapped out the portal to null space as the set of magnitudes of voltage perturbations and the oscillation phases resulting into annihilation of the repeated AP firing. The size of null space portal in the phase-stimulus plot also indicates the size of the null space in the phase space. He also provided experimental evidence that such a null space indeed exists in real neurophysiology of the squid axon, and is not a mere mathematical artifact [29].

However, in real physiological setup, a biological neuron functions by being a part of a network of many interacting neurons in the brain. Further, how the phenomenon of synchronization due to the interacting neurons affects the properties of null space in the phase space is not fully studied. A system of two interacting biological neurons is the most suitable configuration to begin with. Therefore, we apply patterned voltage perturbations to a HH neuron while it is diffusively coupled to another HH neuron and study the null space of the perturbed neurons dynamics. We envisaged that the dynamic coupling currents from another HH neuron might affect null space in the perturbed HH neuron dynamics and is hence worth examining. The coupling interaction between two HH neurons immediately brings forth the property of synchronization between the membrane potential oscillations of the neurons. In fact, there are two prominent kinds of synchronization, i.e. perfectly in-phase and perfectly anti-phase synchronization, which can exist between the repeated firings of action potentials from the coupled HH neurons [30–33]. The present study takes into account both the unique states of synchronization and study the effect of varying coupling strength on the span of null space in the perturbed HH neuron.

The findings demonstrate that the null space exists even in the coupled HH dynamics. Further, the nature of synchronization and the coupling strength between the coupled HH neurons strongly affect the size of null space in the dynamical phase space. Particularly, higher coupling strengths between two in-phase synchronized HH neurons constricts the null space in HH dynamics. However, stronger coupling between two anti-phase synchronized HH neurons expands the null space.

## Methods

### The Hodgkin Huxley model of a neuron

The mathematical description of the dynamics of a HH neuron is given as [1, 2],

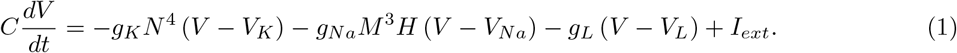

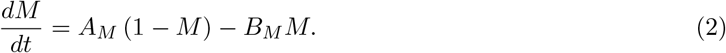

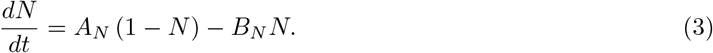

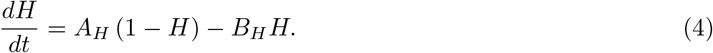

Here,

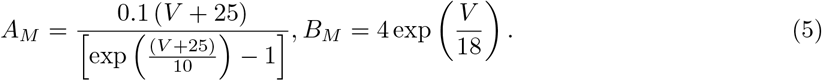

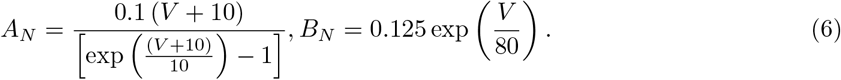

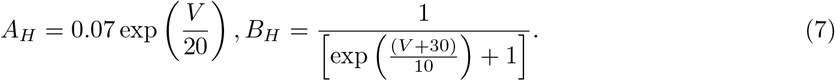

*V* (*t*) describes the membrane potential of the HH neuron at time *t* and is measured in *mV* . *M* (*t*) is the gating variable of the voltage-gated potassium ion (*K*^+^) channels present in the neuronal membrane. *N* (*t*) and *H* (*t*) describe the activation and inactivation, respectively, of the voltage-gated sodium channels. Gating variables express the fractions of active ion channels and, hence, are dimensionless quantities. Owing to its large dimensionality and nonlinearity, an analytical solution of the HH model is unknown. Therefore, the behavior of a HH neuron can be realized only through numerical simulation of the model.

*C* is the capacitance of the neuronal membrane. *g*_*K*_, *g*_*Na*_ and *g*_*L*_ are the conductances of the potassium, sodium and leaky ion channels, respectively. *V*_*K*_, *V*_*Na*_ and *V*_*L*_ are the reversal potentials of the potassium, sodium and leaky ion channels, respectively. *I*_*ext*_ is the constant external stimulating current to the HH neuron. The Equations (5-7) for the *A*s and the *B*s are further sensitive to the temperature *T* at which the neuronal dynamics is observed. These expressions are scaled by a factor of 3^((*T* −6.3)*/*10)^ depending on the temperature of the system. The present study considers the neuronal dynamics to occur at *T* = 6.3^0^*C*, which is the temperature at which Hodgkin and Huxley originally conducted the electrophysiological experiments [1, 2] on the giant squid axon. Best investigated the null space in the HH model using this temperature [29] and we intend to compare the observations on null space in the single isolated HH dynamics with that under coupled situations. Furthermore, electrolyte bath of neuronal cultures and brain slices in many modern electrophysiological studies apply a similar temperature [43]. Accordingly, the Equations (1-7) is directly applicable without any temperature-dependent scaling. In the absence of an external stimulating current, the resting membrane potential of the HH neuron is considered here to be 0*mV* [29]. The magnitudes of the gating variables associated with the resting membrane potential are: *M* = 0.0529, *N* = 0.3177 and *H* = 0.5961. Together, these values of the various dynamical variables under the resting membrane condition define the initial state of the system for numerical simulation. The values of the various parameters are provided in the Supplementary Table.

Further, the behavior of the model critically depends on the magnitude of the constant external current [29]. The vibrant responses of the membrane potential to different values of *I*_*ext*_ are shown in Supplementary Figure 1. For *I*_*ext*_ > 7*µA*.*cm*^−2^ (Supplementary Fig. 1A), there exists a single stable node in the dynamical phase space and the application of current exponentially drives the system to attain the equilibrium state associated with the stable node. For 7*µA*.*cm*^−2^ > *I*_*ext*_ > −6*µA*.*cm*^−2^ (Supplementary Fig. 1B), the existence of stable node transits to that of a stable focus. Consequently, the system is driven by the constant current to the stable focus in a damped oscillatory manner and may once cause appearance of an action potential depending on the initial condition. For −7*µA*.*cm*^−2^ > *I*_*ext*_ > −9*µA*.*cm*^−2^ (Supplementary Fig. 1C), a stable limit cycle appears around the stable focus in the phase space. The limit cycle engenders repeated oscillation in the membrane potential and, thus, leads to the repetitive firing of action potentials under the constant current.

For −10*µA*.*cm*^−2^ > *I*_*ext*_, the existence of the limit cycle continues in the phase space but the stable focus turns into an unstable focus. However, such a change is hardly noticeable in the temporal profile of *V* (*t*) (Supplementary Fig. 1D) as the system still falls into the limit cycle for the given initial conditions of the dynamical variables. Thus, *I*_*ext*_ acts as a critical bifurcation parameter of the model.

For the present study, the *I*_*ext*_ is kept fixed at −8.75*µA*.*cm*^−2^ [29]. The temporal profiles of the various dynamical variables in the presence of chosen *I*_*ext*_ are collectively shown in Figure 1A. The time-period *T*_*P*_ of the oscillation is 15.413*ms* [29]. Simultaneously, the associated stable limit cycle and the stable focus are shown in the two dimensional *V* −*H* phase space (Fig. 1B) as well as the three dimensional *V* −*M* −*N* phase space (Fig. 1C). In the four dimensional *V* −*M* −*N* −*H* phase space, the stable focus is located at the coordinate *V* = −4.96*mV, M* = 0.093, *N* = 0.396 and *H* = 0.42.

**Figure 1.**
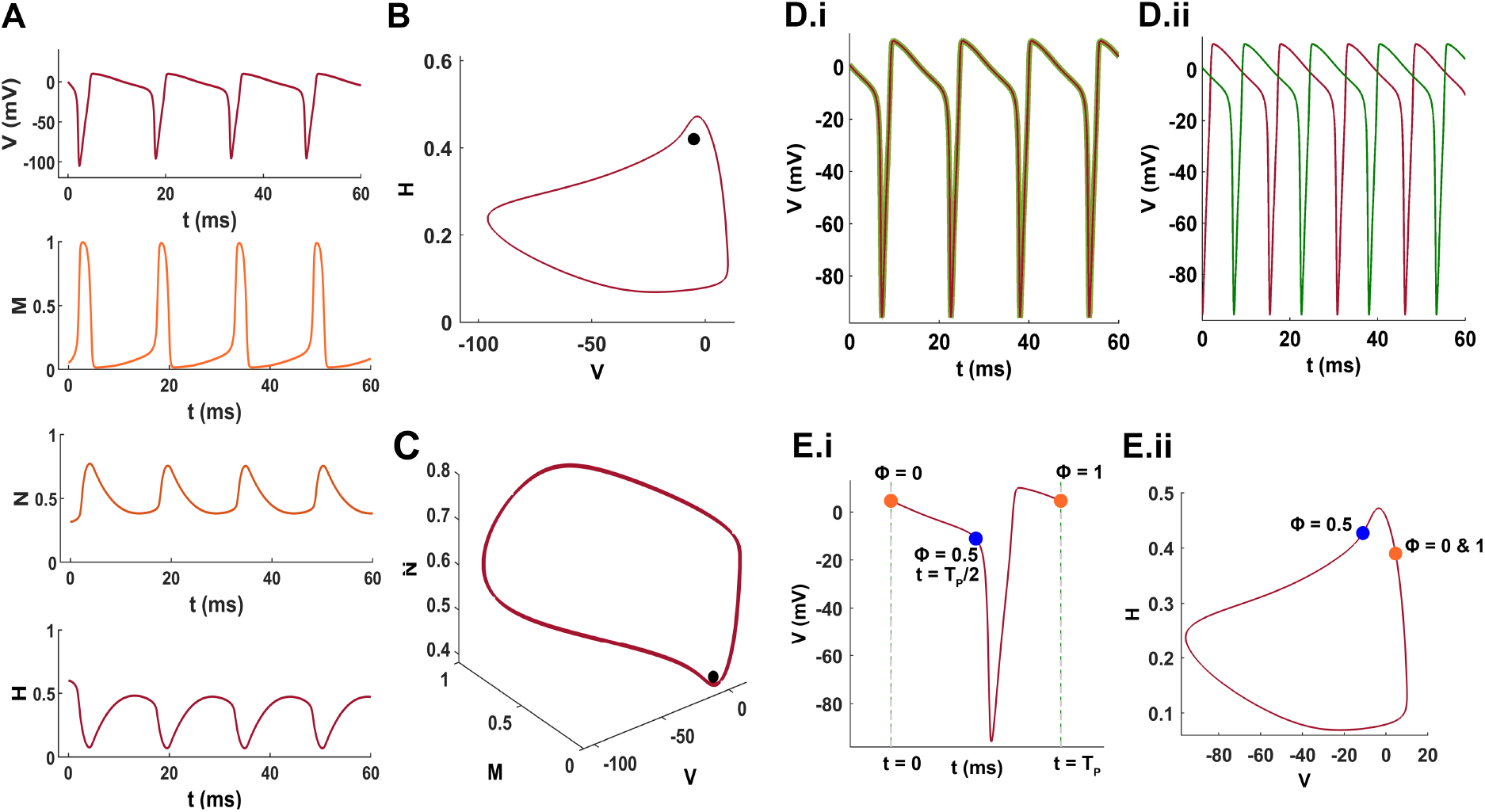
The dynamics of the HH neuron model and the phase synchronization of coupled HH neurons. The external current *I*_*ext*_ is fixed at −8.75*µA*.*cm*^−2^. (A) The membrane potential *V* (*t*) and the various gating variables, *M* (*t*), *N* (*t*) and *H* (*t*), exhibit regular oscillations associated with the repetitive firing of action potentials. (B) For the given initial state, the two dimensional *V* −*H* phase space demonstrates the gradual progression of the system (trajectory in black) towards the limit cycle (shown in red). Notably, there also exists a stable focus (solid green circle) within the limit cycle. (C) The same is illustrated in the three-dimensional *V* −*M* −*N* dynamical phase space of the system. (D.i) Perfectly in-phase synchronized oscillations of two coupled HH neurons with the maximum cross-correlation of 1 completely overlap each other across time. (D.ii) Anti-phase synchronized oscillations are characterized with maximum phase difference and minimum cross-correlation of -0.2259.(E.i) One cycle of membrane potential oscillation with the time-period *T*_*P*_ (15.413*ms*) is shown with the phase points (*ϕ*) ranging [0, 1] linearly distributed over *T*_*P*_ . (*ϕ*) 0, 0.5, and 1 define the initial, middle, and final points, respectively. (E.ii) shows the three phase points on the limit cycle in the *V* −*H* dynamical phase space.

### Coupled dynamics of two HH neurons

The two identical HH neurons are considered to be diffusively coupled. Under the diffusion approximation, the *I*_*coupl*_ to the *i*^*th*^ neuron is taken to be linearly proportional to the difference between the membrane potentials of the coupled neurons (*i* and *j*) and is given by,

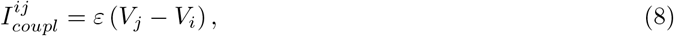

where, the superscript *ij* refers to the direction of *I*_*coupl*_ from *j*^*th*^ neuron to *i*^*th*^ neuron. The proportionality constant *ε* is the coupling coefficient and signifies the strength of diffusive coupling between the two HH neurons. Therefore, by introducing *I*_*coupl*_, the set of ordinary differential equations governing the dynamics of an individual *i*^*th*^ HH neuron is given by,

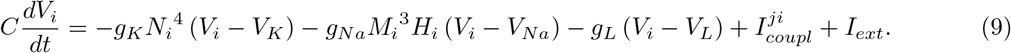

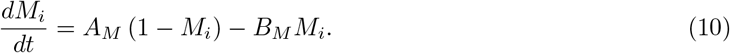

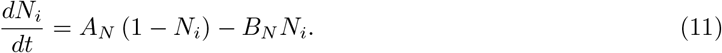

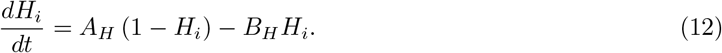

All the parameters of the model are identical for both the neurons. Moreover, the external stimulating current *I*_*ext*_ is identical for both the neurons and is kept fixed at −8.75*µA*.*cm*^−2^, as considered for the single isolated HH neuron.

In the presence of constant *I*_*ext*_, the AP waveforms of the membrane potentials of two coupled HH neurons can exist in synchronization with different magnitudes of phase differences. However, the two most prominent phase relations are the perfect in-phase synchronization and the perfect anti-phase synchronization [30–32]. As the nomenclature suggests, the perfect in-phase synchronization is characterized with zero phase difference between the oscillatory profiles of the neurons (Fig. 1D.i) whereas the perfect anti-phase synchronization represents the maximum phase difference (Fig. 1D.ii). The other possible configurations of synchronization are merely intermediates of these two extreme configurations. In the present study, the perfect in-phase and anti-phase synchronization of the coupled HH neurons are taken into consideration.

### Cross-recurrence plot and its quantitative analysis

To observe the geometry of coupled and synchronized dynamics of HH systems in the four-dimensional phase space, we compute the cross-recurrences between the time-courses of the HH systems [34–36]. Cross-recurrences inform about the time-instances when the two HH systems came in close vicinity of each other while dynamically evolving within the phase space. The state of a single HH system at any time, *t*, in the phase space is defined by 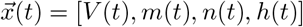. Therefore, we compute the pair-wise Euclidean distance (∥ . ∥) between the coupled HH state vectors 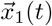 and 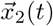 at their respective time-points *t*_*i*_ and *t*_*j*_ as,

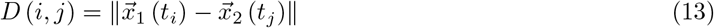

By choosing a scalar cut-off value *δ*, we can define close vicinity of the state vectors 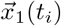 and 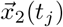 in the phase space if the *D* (*i, j*) ≤ *δ*. Therefore, the cross-recurrence, *R*(*t*_*i*_, *t*_*j*_), is given as [34, 35],

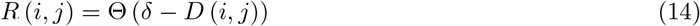

Θ(.) represents the heavy-side step function. *R*(*t*_*i*_, *t*_*j*_) is 1 under the arbitrarily defined close vicinity, otherwise is 0. Computing *D*(*t*_*i*_, *t*_*j*_) and *R*(*t*_*i*_, *t*_*j*_) for all pairs of *t*_*i*_ and *t*_*j*_ results into a square distance matrix, *D*, and cross-recurrence matrix, *R*, respectively, of size *N* × *N*, where *N* is the number of uniform discrete time-points in the time-series 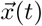. Plotting R provides cross-recurrence plot (CRP), which visually summarizes the geometry and stability of synchronized dynamics, the manifolds of slow-fast timescales, and intermittent epochs of de-synchronization, if any.

Next, we perform a quantitative analysis of the CRP by computing recurrence rate (RR) and determinism (DET). The RR represents the magnitude of recurrence and is described as the fraction of *N* × *N* pairs of time-points when the two HH systems are in close vicinity in the phase space [37],

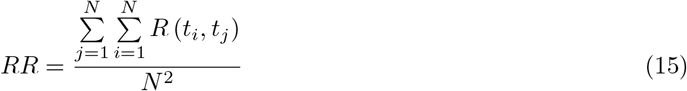

Apparently, *δ* strongly affects the value of RR. Based on literature [38], we choose a *RR* = 0.10 to compute CRP consistently for all cases of synchronization studied here. Accordingly, we calculate *δ* as the 10th percentile of the distribution of Euclidean distance in the matrix *D*. Further, *RR* can also be computed along the diagonals of the matrix *R*. Since the diagonals in *R* inform about the time lags, plotting the diagonal *RR*s, denoted here as *RR*_*D*_, can provide a more straightforward picture of the phase shifts and pattern repetitions under different states of synchronization.

Determinism (*DET*) defines the fraction of recurrence points that form diagonal lengths greater than or equal to a minimum length (≥ *l*_min_) and is given as [39, 40],

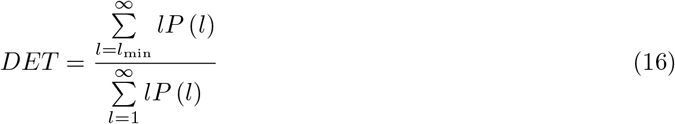

≥ *l*_min_ = 2 is typically chosen as the minimum length criteria [41]. *DET* informs about the determinism in the synchronized dynamics, where lower *DET* values closer to 0 underscore randomness in coupling dynamics and higher *DET* close to 1 reflects more deterministic interactions between the coupled HH systems.

### Voltage perturbation and Phase Response Curve (PRC)

When the HH neuron is repeatedly producing APs under constant *I*_*ext*_ (−8.75*µA*.*cm*^−2^), a voltage perturbation of strength *dV* is given to the neuron by adding *dV* to the neurons membrane potential *V* (*t*), i.e. *V* (*t*) + *dV*, at a particular instance of time, *t* [29]. *dV* can be a positive or a negative magnitude of the perturbation. The desired time-points of voltage perturbations are decided in terms of phase points in one cycle of oscillation associated with the AP. However, it solely depends on how an observer arbitrarily chooses to map one cycle of oscillation on the temporal profile. The choice of mapping will not affect the quality of voltage perturbation study, except that the perturbation-induced features will be merely shifted along the quantitatively different descriptions of phase points. As the features of a single HH neuron studied by Best [29] is the benchmark for the present comparative investigation, the initial point of a cycle of oscillation is chosen as the systems state given by *V* = 4.82*mV, M* = 0.030, *N* = 0.450 and *H* = 0.397, to maintain consistency.

Figure 1E.i shows the one cycle of oscillation in terms of the temporal profile of the membrane potential and Figure 1E.ii shows the same through the corresponding limit cycle in the two-dimensional *V* −*H* phase space. Given the initial point, the phase points (*ϕ*) of the oscillation cycle are described in a range [0, 1], where *ϕ* = 0 corresponds to the beginning of the oscillation cycle at a reference origin time *t* = 0 and *ϕ* = 1 corresponds to the end of the oscillation at time *t* = *T*_*P*_ . Accordingly, *ϕ* is assigned a value as the fraction of *T*_*P*_ which, in turn, signifies the fraction of cycle completed along the time. For instance, *ϕ* = 0, 0.5 and 1.0 corresponds to the 0%, 50% and 100% completion of the cycle, respectively, with respect to the time-period. In fact, such an assignment of phase points in the oscillation cycle has a linear distribution over the time-period of the cycle.

In the case of two coupled HH neurons, the voltage perturbation is given to only one of the neurons. This is to satisfy the present prime objective to comparatively enquire the nature of perturbations in a HH neuron when it is isolated and when it is coupled to another HH neuron. Undoubtedly, paired-perturbations in both the neurons is an interest of future investigation. Here, the neuron #1 (*i* = 1) is chosen for the voltage perturbation throughout the study.

After applying a voltage perturbation of strength *dV* at a given phase point *ϕ* of the oscillation cycle, the response of the perturbed HH neuron is examined after a time-duration equivalent to *n* multiples of the time-period *T*_*P*_ [25, 26]. The phase of oscillation after a voltage perturbation is termed as the new-phase [29] and is here denoted by *φ*. Corollary, the phase *ϕ* at which the voltage perturbation is applied becomes the old-phase (*ϕ*) [29]. Here, the analysis consistently uses *n* = 20, so that enough time is assured for the return of the oscillator to its limit cycle after a voltage perturbation. Thus, a PRC is obtained between old and new phases for a perturbation *dV* [25, 26, 29, 33]. There are classically two kinds of PRCs ubiquitously observed in almost all periodic natural systems: type 1 and type 0 [25, 29]. Type 1 PRCs are characterized by new phases shifted away from the old phases by almost a constant amount. Visibly, *φ* monotonously increases along *ϕ* and covers the cycle of phases 0 to 1. This leads to an average PRC slope close to 1. However, type 0 PRCs are characterized by new phases with inconsistent shifts from the old phases. *φ* exhibits non-monotonous behaviour and can also undergo sudden jumps in its value along *ϕ*. This leads to an average PRC slope close to 0.

However, in the bistable regime, it is not always that the HH neuron may return to its limit cycle after a perturbation. Certain perturbations may also lead to the attraction of the system to the stable focus and the annihilation of oscillation associated with the repetitive AP firing. In this situation, the new phase is impossible to be defined and, hence, is referred to as phase-indeterminate [25, 29]. Typically, voltage perturbations leading to phase-indeterminate outcomes for a stretch of old phases *ϕ* are also characterized by Type 0 class, as the new phases *φ* at the boundaries of phase-indeterminate window exhibit non-monotonous trend [25, 29].

The scripts for the numerical simulations of the HH dynamical equations were written in MATLAB R2018b (The MathWorks) and the built-in ode15 solver was used for the numerical integration of the differential equations. The scripts will be made publicly available after the publication of the manuscript.

## Results

### Phase resetting behavior of a single HH neuron

A voltage perturbation can lead to two major outcomes: the perturbed HH neuron either regains its oscillation of the membrane potential underlying the repetitive firing of action potentials or looses oscillation and the repetitive firing is annihilated. Figure 2 demonstrate these outcomes under the applications of different strengths of voltage perturbation at an identical phase point and identical strength of perturbation at different phase points, respectively. In the *V* −*H* phase space, this becomes more clear as the system post-perturbation either returns to the stable limit cycle or falls in the trap of stable steady state (Fig. 2). These observations convey an important message that neither the strength nor the phase point of the voltage perturbation alone is crucial in deciding the fate of the perturbed system.

**Figure 2.**
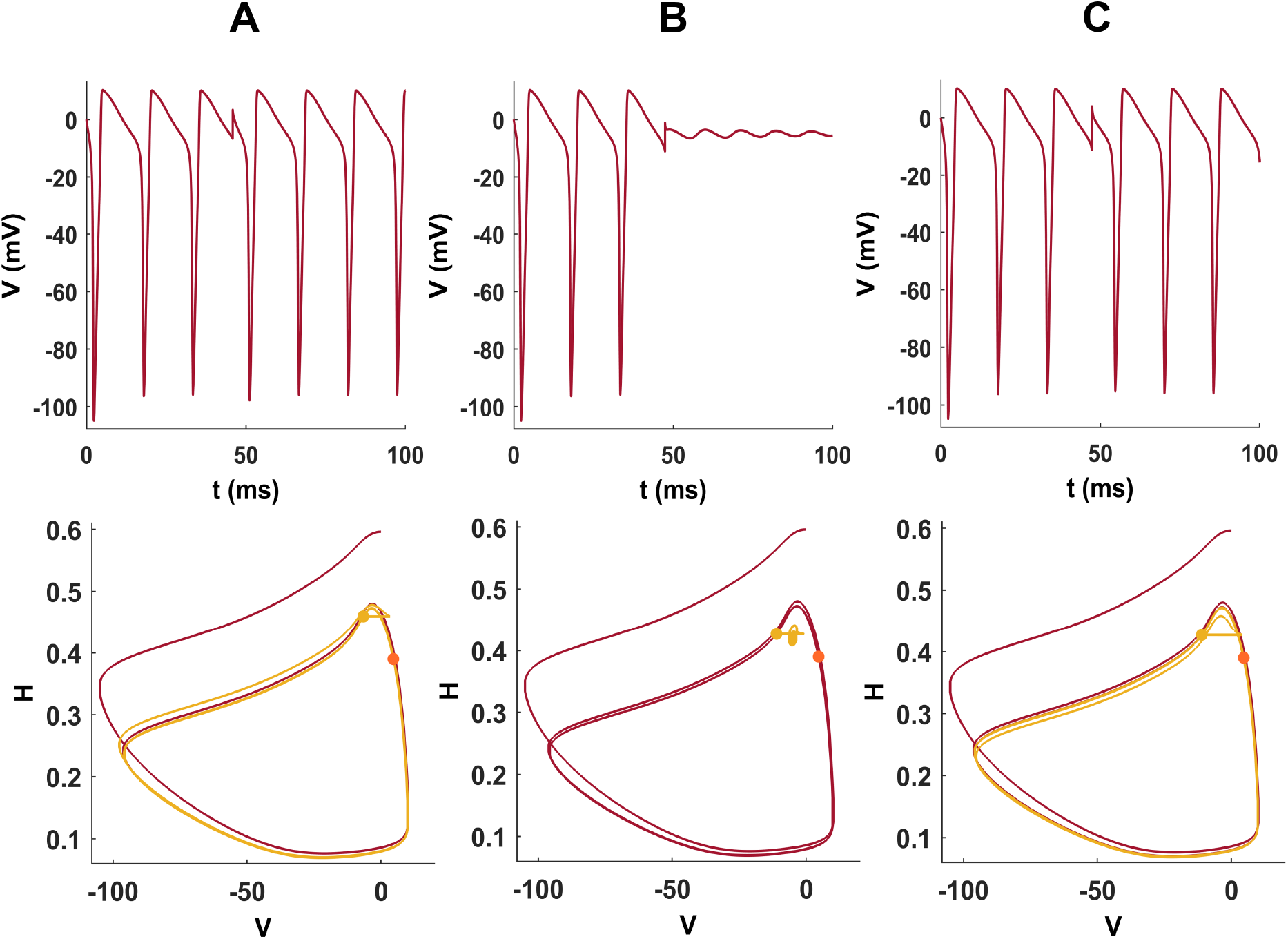
Effects of voltage perturbations on the membrane potential oscillation in an isolated HH neuron. (A) Top panel; A voltage perturbation of strength *dV* = 10*mV* at the phase point *ϕ* = 0.4 leads to restoration of the repeated membrane potential oscillation. Bottom panel; The same is shown in the *V* −*H* dynamical phase space, where the post-perturbation system trajectory (shown in light blue) gradually returns to the original limit cycle of the oscillation (in red).(B) Same strength of the voltage perturbation *dV* = 10*mV* but at the phase point *ϕ* = 0.5 causes the annihilation of the repeated membrane potential oscillation (top panel) with the post-perturbation trajectory collapsing into the stable focus. (C) However, a slightly higher perturbation strength of *dV* = 11*mV* at the same phase point *ϕ* = 0.5 leads to restoration of the repeated membrane potential oscillation (top panel) with the post-perturbation trajectory returning to the original limit cycle (bottom panel).

The restoration of the oscillation in membrane potential after a voltage perturbation may further be associated with a difference in the old phase (*ϕ*) of the perturbation and the new phase (*φ*) of the restored oscillation. This is effectively captured in the PRCs for different perturbation strengths (Fig. 3). For positive perturbation strengths, a small magnitude *dV* = 2*mV* yields a type 1 PRC (Fig. 3A.i), whereas stronger perturbations (*dV* = 5, 10, &60*mV* ) results into type 0 PRCs (Fig. 3A. ii, iii, & iv). An important thing to note is the range of old phases around *ϕ* = 0.5 which leads to phase indeterminate situation resulting from the annihilation of repeated membrane potential oscillation to the stable steady state. These phase indeterminate regions depict the portal to the null space, where perturbations can throw the system away from the basin of attraction associated with the limit cycle in the dynamical phase space. Similarly, for the negative values of voltage perturbations, a small magnitude *dV* = −2*mV* yields a type 1 PRC (Fig. 3B.i), whereas stronger perturbations (*dV* = −5, −10, *&* − 60*mV* ) results into type 0 PRCs (Fig. 3B. ii, iii, & iv). Here, the range of old phases around *ϕ* = 0.2 leads to phase indeterminate situation. The transition of PRC from type 1 to type 0 is a key factor to be observed, as it signifies the critical intensity of voltage perturbation which brings the neuronal dynamics in close vicinity of the null space associated with the basin of attraction of the stable focus. In the case of single HH neuron, the transition of PRC from type 1 to type 0 happens at *dV* = 4*mV* and *dV* = −4*mV* along the increasing intensity of the positive and negative voltage perturbations, respectively.

**Figure 3.**
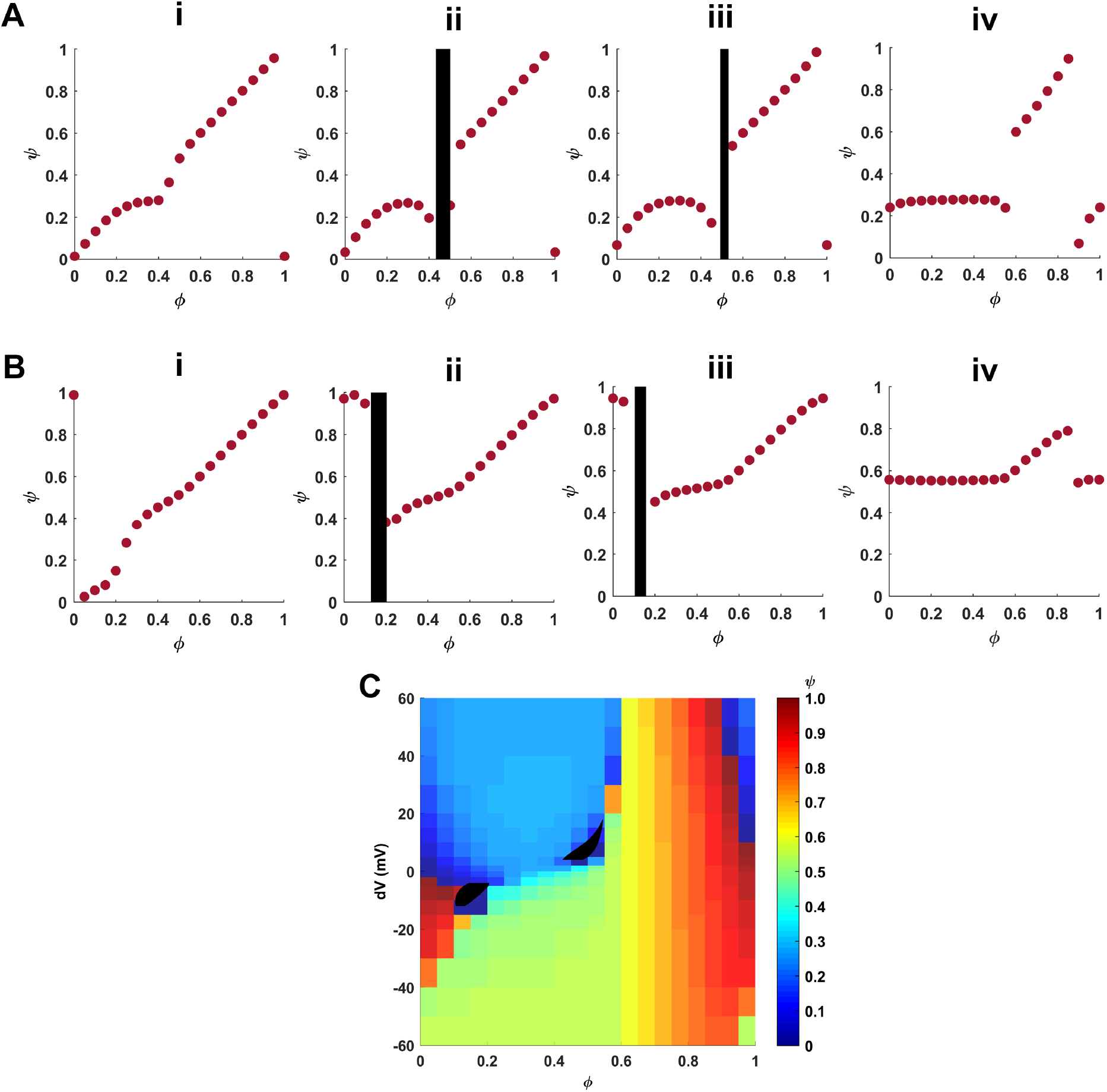
Phase response curves (PRCs) of a single HH neuron for different voltage perturbations and the phase-stimulus map. PRC describes the relation between the new phase points (*φ*) at which the HH system re-enters the limit cycle after a voltage perturbation of certain strength is applied at the old phase points (*ϕ*) of the originally oscillating system on the limit cycle. Those *ϕ* at which the voltage perturbations lead to annihilation of the repeated oscillation accounts for phase indeterminate situations, where the system enters the stable focus within the limit cycle. (A) For positive voltage perturbations, *dV* = 2*mV* (i) results into type 1 PRC whereas *dV* = 5*mV* (ii), 10*mV* (iii), and 60*mV* (iv) leads to type 0 PRCs. The vertical black bars in (ii) and (iii) depict the phase indeterminate situation. (B) For negative voltage perturbations, *dV* = −2*mV* (i) results into type 1 PRC whereas *dV* = −5*mV* (ii), −10*mV* (iii), and −60*mV* (iv) leads to type 0 PRCs. The vertical black bars in (ii) and (iii) depict the phase indeterminate situation. (C) Based on PRCs, a phase-stimulus plot is obtained where, for each perturbation strength (*dV* ) and old phase point (*ϕ*), the corresponding new phase (*φ*) is shown using the color scale (on the right). The plot also serves as an isochron map to identify the regions of identical new phases. The set of perturbation strengths and old phases which together lead to phase indeterminate situations or annihilation of oscillation is shown as a bounded black patch on the map. This region serves as a portal to the null space or the basin of attraction of the stable focus. Here, the values of the *dV* are ±[0, 2, 5, 10, 15, 20, 30, 40, 50, 60]*mV* and *ϕ* is discretized with a step of 0.05. The overlaid black patch of null space portal has been created with finely discretized values of *dV* and *ϕ* with step-sizes of 1*mV* and 0.001, respectively.

Using the PRCs acquired for many more positive and negative values of *dV*, a phase-stimulus plot is obtained (Fig. 3C), which also serves as an isochron map. The term isochron refers to the set of different strengths and old phases of voltage perturbation resulting into an identical new phase. The isochron map helps in tracking the robust phase points of the oscillation in membrane potential. Here, the range of *ϕ* = 0.6−0.8 appears to be quite robust phase points, with *ϕ* = 0.6 as the most robust phase. Perturbations of any strength applied at these old phases lead to restoration of the oscillation at new phases nearly identical to the old phases.

The old phases associated with the phase-indeterminate situation across varying *dV* collectively constitute the subset of *dV* and *ϕ* which leads to the collapse of perturbed neuronal dynamics into the limit cycle’s null space, which is also the basin of attraction of the stable focus. The phase-indeterminate *dV* and *ϕ* sets are shown as black patches in the phase-stimulus plot (Fig. 3C) and are commonly referred to as the portal to the null space [24, 29]. The size of the null space is a crucial indicative measure of the spatial span of the steady state’s basin in the dynamical phase space, such that larger null space indicates larger span of the basin. Moreover, both very low and very high perturbation strengths, regardless of the sign, skip the null space. Only a moderate range of perturbation strengths can cause the system to enter the null space. For positive perturbations, (*dV* = 4−18*mV* applied at *ϕ* within [0.4−0.55] pushes the system into the null space. For negative perturbations, entry to null space occurs for *dV* = −4− − 12*mV* applied at *ϕ* within [0.1−0.25]. The two null space portals are almost symmetrically located around *ϕ* = 0.3, which is also understandable given the time period-based distribution of phase values on the stretch of limit cycle in close vicinity of the stable focus (Fig. 1B & E) and given that only the voltage perturbation is being performed.

These results obtained for the single HH neuron are identical to that observed in the voltage perturbation study by Best [29]. Further, these observations would serve as the control or benchmark for the comparative study of voltage perturbations in the coupled HH neurons.

### Phase resetting behavior of a pair of coupled perfectly in-phase synchronized HH neurons

We consider a diffusively-coupled system of two identical HH neurons firing repeated APs in perfect in-phase synchronization. We vary the coupling coefficient *ε* over orders of magnitude, *ε* ∈ [1*e* − 4, 1*e* − 3, 5*e* − 3, 1*e* − 2, 2.5*e* − 2, 5*e* − 2, 7.5*e* − 2, 1*e* − 1]. A sufficiently weak coupling between two interacting systems is a pre-requisite for synchronization [30]. In the present setup, the diffusive coupling current is proportional to the difference of neurons’ membrane potentials, which can potentially reach to orders of 10 during perturbations. Therefore, we limit *ε* to a maximum value of 0.1 [31]. The limit cycles of the two coupled HH neurons and their time periods remain identical to that of the single isolated HH neuron.

The cross-recurrence plot (CRP) using 60*ms*-long time-series data of the coupled HH systems for *ε* = 1*e* − 4 (Fig. 4A, left panel) shows the recurrence structure lying along the main diagonal, and the structure is repeated off-diagonally at a regular gap of the time-period (*T*_*p*_ = 15.413*ms*) of the AP waveform. The violin-shaped pattern in the recurrence structure underscores the nonlinear slow and fast timescales of the AP waveform. The diagonal-wise recurrence rate (*RR*_*D*_) plot (Fig. 4A, right panel) clearly shows the *RR*_*D*_ peak exactly at the zero time-lag and repetition of the peak at the time-lags equal to integer multiples of *T*_*p*_. The peak *RR*_*D*_ value very close to 1 informs almost continuous recurrent structure with less break points along the diagonals. The CRP and *RR*_*D*_ plots remain identical for all values of *ε*. The determinism (*DET* ) is also identically 1 across all values of *ε*. These features demonstrate robust and deterministic in-phase synchronization between the coupled HH systems.

**Figure 4.**
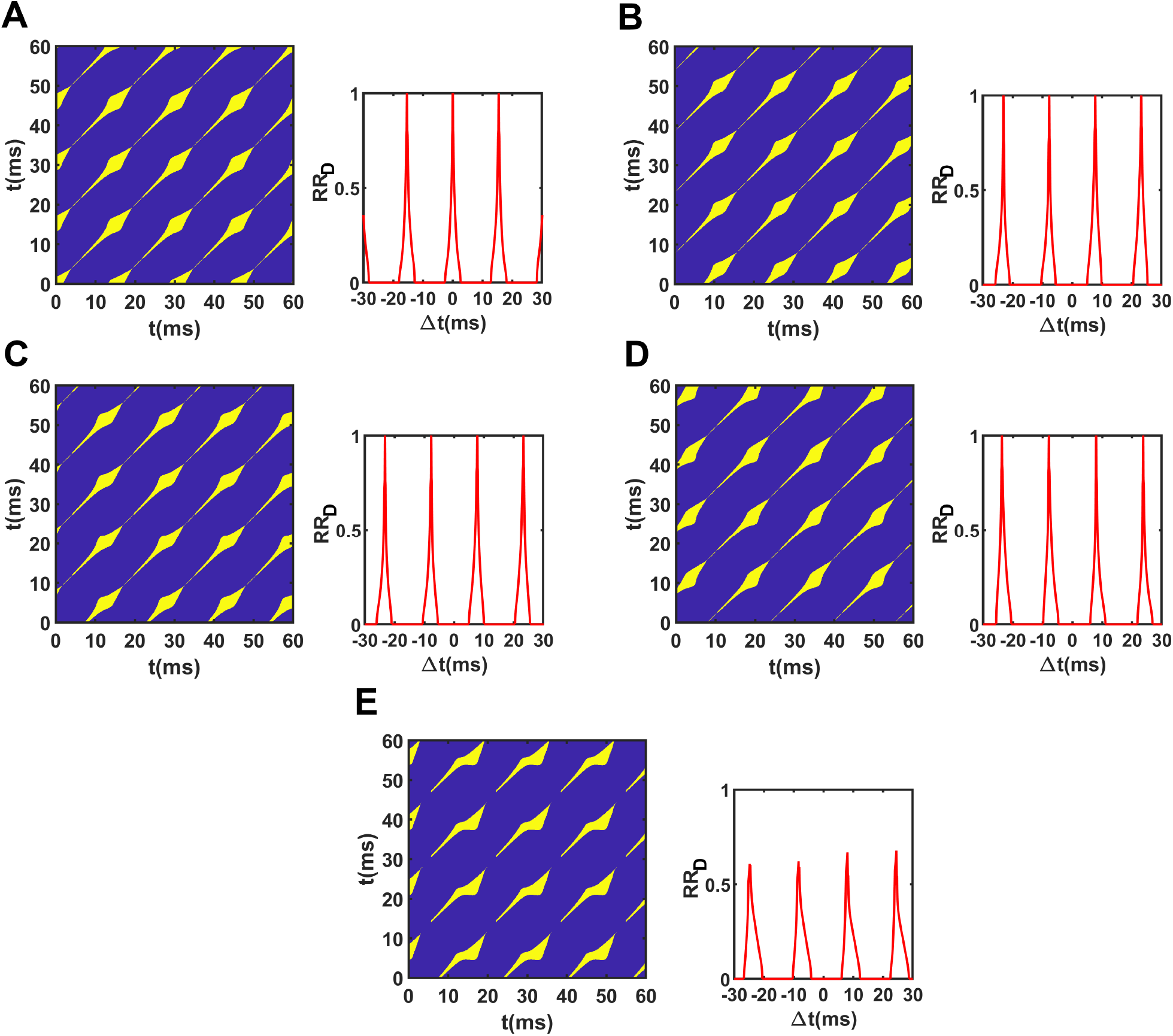
Cross-recurrence plot (CRP) and diagonal-wise recurrence rates (*RR*_*D*_) of perfect in-phase and anti-phase synchronized coupled HH dynamics. (A) The CRP (left panel) of the 60*ms*-long coupled HH dynamics under perfect in-phase synchronization with coupling coefficient *ε* = 1*e* − 4 shows recurrence structure (shown in yellow) exactly along the main diagonal line. The recurrence structure appears further repeated in parallel off-diagonally at a regular gap identical to the time-period (*T*_*p*_ = 15.413*ms*) of the AP waveform. The violin-shaped expanding and contracting design of the recurrence structure arises from the nonlinear slow and fast timescales of the HH dynamics. The plot of *RR*_*D*_ (right panel) demonstrates the peaks of *RR*_*D*_ at zero time-lag(*δt*) as well as at time-lags equal to integer multiples of *T*_*p*_. The peak *RR*_*D*_ value approaching 1 indicates almost continuous recurrence structure at the corresponding time-lags. (B-E) The CRP plots (left panel) under perfect anti-phase synchronization with *ε* = 1*e* − 4(B), 1*e* − 3(C), 5*e* − 3(D) and 1*e* − 2(E), consistently show the off-set of recurrence structure from the main diagonal. This off-set is identical to half of the *T*_*p*_ and confirms perfect anti-phase synchronization across different values of *ε*. This becomes more evident in the *RR*_*D*_ plots (B-E, right panels) for the different values of *ε*. The peak *RR*_*D*_ is shifted away from the origin by a time-lag equal to the half of *T*_*p*_. Moreover, the *RR*_*D*_ peaks are repeated on either sides at a regular gap of *T*_*p*_ further to the original off-set. The small *RR*_*D*_ peak values (much less than 1) for *ε* = 1*e* − 2 (E, right panel) indicate highly fragmented recurrence structure, as visible in the associated CRP (E, left panel).

Voltage perturbations are applied to only one of the neurons. The two possible outcomes of the voltage perturbations, either restoration or annihilation of membrane potential oscillation (Supplementary Fig. 2) shows that the perturbed neuron has negligible effect on the unperturbed neuron. This remains true for all *ε*. Further, if the perturbed neuron’s oscillation gets restored, the initial perfect in-phase synchronization continues to exist.

For each *ε*, PRCs of the perturbed neuron are obtained for positive as well as negative values of voltage perturbations. For *ε* = 1*e* − 4, 1*e* − 3, 1*e* − 2&1*e* − 1, PRCs of the perturbation magnitudes 2, 5, 10, &60*mV* are shown in Supplementary Figure 3, 4, 5, & 6, respectively. Similar to the previous observations on a single isolated HH neuron, sufficiently weak (for instance, *dV* = ±2*mV*, Supplementary Fig. 3) and strong perturbations (*dV* = ±60*mV*, Supplementary Fig. 6) lead to type 1 and type 0 PRCs, respectively, for all values of *ε*. The critical positive voltage perturbation causing transition from type 1 to type 0 PRC for *ε* = 1*e* − 4 is 3*mV*, which is slightly lower than that (4*mV* ) observed for the single HH neuron. For *ε* = 1*e* − 3&1*e* − 2, the critical positive voltage perturbation is 4*mV* . However, for *ε* = 1*e* − 1, the critical value shifts to very high value, 18*mV* . The critical negative voltage perturbation causing transition from type 1 to type 0 PRC for *ε* = 1*e* − 4, 1*e* − 3, &1*e* − 2 is −4*mV*, identical to that (−4*mV* ) observed for the single HH neuron. However, for *ε* = 1*e* − 1, the critical negative perturbation value jumps to -14mV.

Using PRC data, the phase-stimulus maps for the different coupling coefficients are obtained. Figure 5A shows phase-stimulus plots for *ε* = 1*e* − 4, 1*e* − 3, 1*e* − 2&1*e* − 1. Similar to the single isolated HH neuron, the range of *ϕ* = 0.6−0.8 appears to be quite robust phase points, with *ϕ* = 0.6 as the most robust phase. Another interesting thing to note is the variation in the shape and size of the null space portals in the positive as well as negative regions of voltage perturbations across the coupling coefficients (Fig. 5A). To obtain a quantitative estimate of the area of a null space portal in the phase-stimulus plot, we summed together the lengths of the phase-indeterminate stretches of the old phases for the individual perturbation magnitudes distanced by unit step (1*mV* ). Here, we assume that the stretch of phase-indeterminate old phases remains constant over the unit interval of perturbation magnitude and discretize the patch of null space portal into rectangular grids of size *lengthsofthephase* − *indeterminatestretch* × 1*mV* . The areas of the null space portals in the positive and negative regions of voltage perturbations are separately computed. In comparison to the single isolated HH neuron (Fig. 5B.i), the null space portal for positive perturbations initially expands with increase in coupling coefficient. However, a sharp reduction in the portal area is observed around *ε* = 0.01 and later vanishes for higher coupling coefficients. On the contrary, the size of null space portal for negative perturbations (Fig. 5B.ii) monotonously decrease with increase in coupling coefficient and vanishes to almost zero for very high coupling strength. Altogether, the decreasing trend of the total size of the null space portal (Fig. 5B.iii) indicates a restricted passage of the voltage-perturbed repeatedly oscillating system to the null space of its limit cycle for higher coupling strengths between two in-phase synchronized HH neurons.

**Figure 5.**
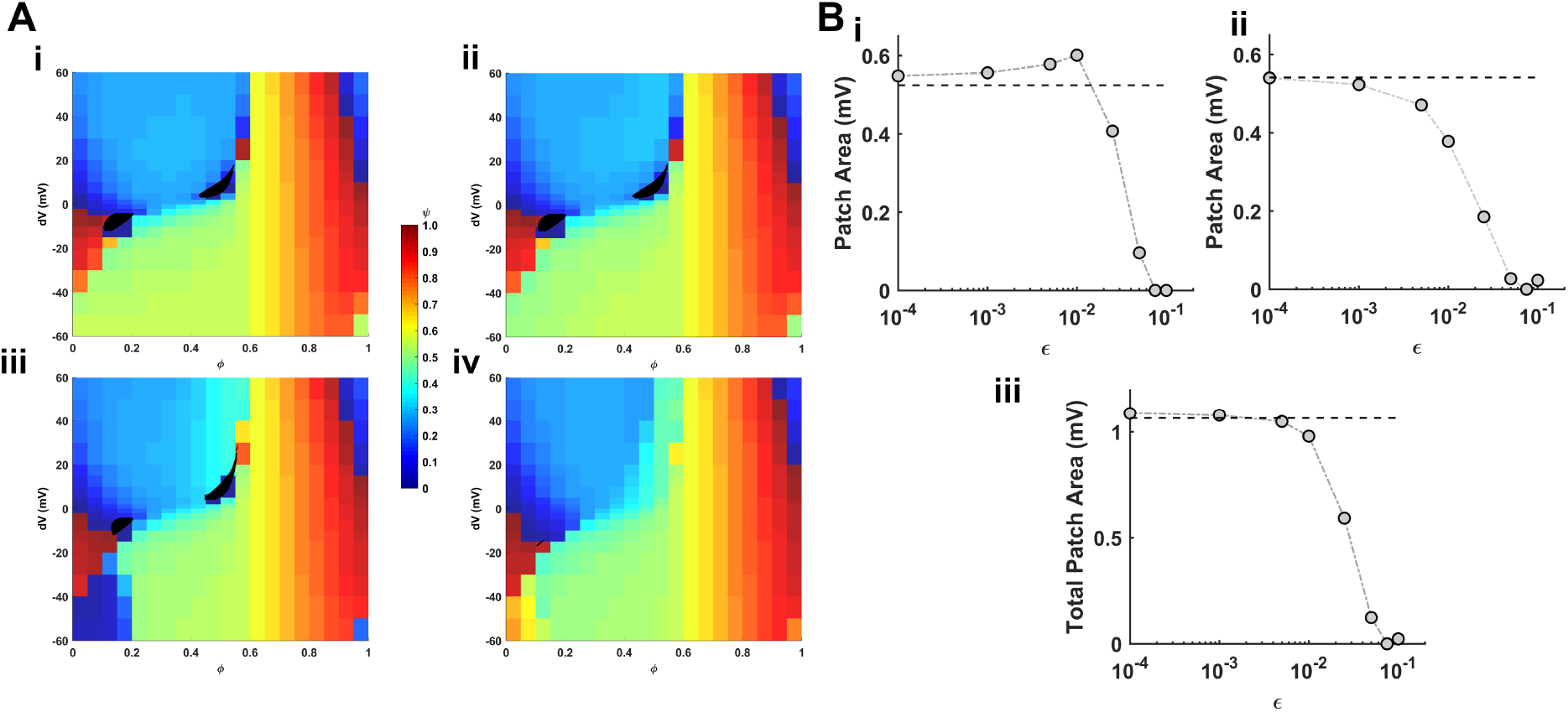
Phase-stimulus plots and null space portals for different coupling coefficients of two in-phase synchronized HH neurons. (A) For the coupling coefficients *ε* = 1*e* − 4(i), 1*e* − 3(ii), 1*e* − 2(iii) and 1*e* − 1(iv), phase-stimulus plots show new phases (*φ*) of the restoration of oscillation, using the color scale (on the right), when one of the coupled HH neurons is perturbed with different perturbation strengths (*dV* ) at the old phase points (*ϕ*) of the limit cycle. These plots also serve as isochron maps where voltage perturbations leading to identical new phase values can be identified based on identical colour. The set of perturbation strengths and old phases which together lead to phase indeterminate situations or annihilation of oscillation is shown as a bounded black patch on the map. This region serves as a portal to the null space or the basin of attraction of the stable focus. Here, the values of the *dV* are ±[0, 2, 5, 10, 15, 20, 30, 40, 50, 60]*mV* and *ϕ* is discretized with a step of 0.05. The overlaid black patch of null space portal has been created with finely discretized values of *dV* and *ϕ* with step-sizes of 1*mV* and 0.001, respectively. (B) Variation in the area of null space portals in the regions of positive (i) and negative (ii) perturbations of the phase-stimulus plot across increasing coupling coefficient *ε*. The dashed black line shows the area of null space portal for a single isolated HH neuron. The area of the null space portal is computed by adding together the spans of phase indeterminate *ϕ* over *dV* discretized by the step-size of 1mV. Variation in the total area of null space portal (iii) in the phase-stimulus plot across increasing *ε*.

### Phase resetting behavior of a pair of coupled perfectly anti-phase synchronized HH neurons

Next, we consider a diffusively-coupled system of two identical HH neurons firing repeated APs in perfect anti-phase synchronization. Again, voltage perturbations are applied to only one of the neurons. The value of coupling coefficient *ε* is chosen over orders of magnitude, *ε* ∈ [1*e* − 4, 1*e* − 3, 2.5*e* − 3, 5*e* − 3, &1*e* − 2]. Note that the maximum (*ε* = 1*e* − 2), as opposed to 1*e* − 1 previously. A theoretical investigation by Ao et al. [42] of the nature of synchronization in HH neurons showed that the anti-phase synchronization is not a very robust configuration and the maximum coupling coefficient suitable for a stable anti-phase synchronization is *ε* = 0.01. For higher *ε*, the anti-phase configuration becomes extremely unstable and rapidly collapses into a state of perfect in-phase synchronization (Supplementary Fig. 7A).

Moreover, a perfect anti-phase synchronization stretches the time-periods of the oscillation of the individual HH neurons identically, compared to that for a single HH neuron. Higher coupling coefficient leads to longer time period (Supplementary Figure 7B.i). However, the shape and size of the limit cycle in the phase space remains identical to that for a single HH neuron across all coupling coefficients (Supplementary Figure 7B.ii). Change in the time-period also necessitates linear reallocation of the phase points [0, 1] over the limit cycle for different coupling coefficients.

The CRPs using 60*ms*-long time-series data of the coupled HH systems for *ε* = 1*e* − 4, 1*e* − 3, 5*e* − 3, and 1*e* − 2 (Fig. 4B-E, left panels) show offset of the recurrence pattern from the main diagonal. This off-set is identical to half of the *T*_*p*_ of the AP waveform, which is desired owing to the perfect anti-phase synchronization. Following the offset from the main diagonal, the recurrence pattern is repeated off-diagonally at a regular gap of the *T*_*p*_. Here again, the violin-shaped pattern underscores the nonlinear slow and fast timescales of the AP waveform. The diagonal-wise recurrence rate (RR) plots (Fig. 4B-E, right panels) clearly show the RR peaks at time-lag identical to half of the *T*_*p*_ and is further repeated on either sides at a gap of *T*_*p*_. The peak *RR* very close to 1 (Fig. 4B-D, right panels) informs almost continuous recurrence, whereas short *RR* peaks for *ε* = 1*e* − 2 (Fig. 4E, right panel) reflects the broken pattern along diagonals evident in the corresponding CRP (Fig. 4E, left panel). The determinism (*DET* ) is identically 1 across all values of *ε* and demonstrates deterministic as well as robust perfect anti-phase synchronization between the coupled HH systems.

The two potential outcomes, restoration and annihilation of the oscillation, due to voltage perturbations in the perturbed neuron are shown in Supplementary Figure 8. Similar to the case of in-phase synchronization, the perturbations in one neuron has negligible effect on the unperturbed neuron. Further, if the perturbed neuron’s oscillation gets restored, the initial perfect anti-phase synchronization collapses to perfect in-phase synchronization post-perturbation. This is again due to the fragile nature of the anti-phase synchronization in HH systems.

For each *ε*, PRCs of the perturbed neuron are obtained for positive as well as negative values of voltage perturbations. For *ε* = 1*e* − 4, 1*e* − 3, 5*e* − 3, &1*e* − 2, PRCs of the perturbation magnitudes 2, 5, 10&60*mV* are shown in Supplementary Figure 9, 10, 11, & 12, respectively. Contrary to the situations of single HH neuron and in-phase synchronized HH neurons, very weak perturbation of 2*mV* results into type 1 PRC only for sufficiently low coupling strengths (Supplementary Fig. 9) and leads to type 0 PRC for high coupling strength. The higher perturbation strengths (Supplementary Fig. 10, 11, & 12) consistently yield type 0 PRCs. However, compared to previous situations, type 0 PRCs appear with large irregular dispersion of the new phases with higher irregularity for higher coupling coefficient. The critical positive perturbation strengths causing transition from type 1 to type 0 PRC are 3*mV* for *ε* = 1*e* − 4, 1*e* − 3, &5*e* − 3 and 2*mV* for *ε* = 1*e* − 2. The critical negative perturbation strengths causing transition from type 1 to type 0 PRC are −4*mV* for *ε* = 1*e* − 4&1*e* − 3 and −3*mV* for *ε* = 5*e* − 3&1*e* − 2.

The phase-stimulus plots for *ε* = 1*e* − 4, 1*e* − 3, 5*e* − 3, &1*e* − 2 (Fig. 6A) shows that the range of old phase *ϕ* = 0.6−0.8 is robust against perturbation only for lower values (*ε* = 1*e* − 4&1*e* − 3) of the coupling coefficient. However, for higher (*ε* = 5*e* − 3&1*e* − 2) coupling coefficients, these phases loose their robustness. The old phase *ϕ* = 0.6 remains most robust for all values of coupling coefficients. Moreover, higher (*ε* = 5*e* − 3&1*e* − 2) coupling coefficients exhibit apparently high irregularity in the isochron map. Another striking thing to note is the presence of grey patches on the phase-stimulus plots (Figure 6A) for higher values of coupling coefficients *ε* = 5*e* − 3&1*e* − 2. The color choice is opposed to the familiar black patches representing null space portals. The grey patches belong to a combination of perturbation strengths and old phases which neither annihilate nor restore the original oscillation of the perturbed neuron. Rather, the system enters a new limit cycle with phase space span much smaller than the original limit cycle. Interestingly, the new extremely small limit cycle remains seated closely around the stable focus. The size of null space portals also strongly vary with increase in the coupling coefficient (Fig. 6A). The areal span of the null space portals, in the positive (Fig. 6B.i) as well as negative (Fig. 6B.ii) regions of the voltage perturbations, consistently increase with increase in the coupling coefficient. Therefore, the net size of the null space portal (Fig. 6B.iii) also monotonously increases and is higher than that of the single HH neuron as well as of the in-phase synchronized HH neuron pairs.

**Figure 6.**
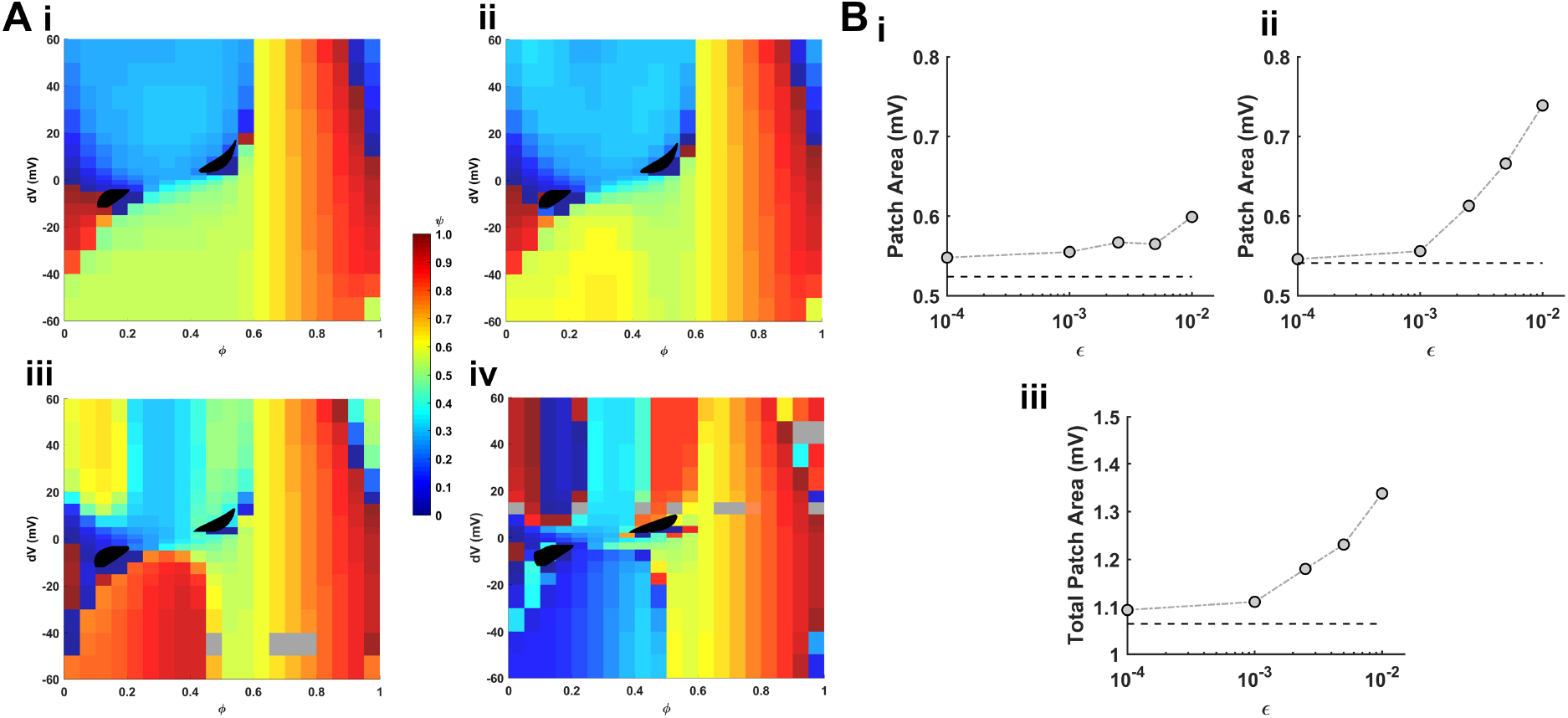
Phase-stimulus plots and null space portals for different coupling coefficients of two anti-phase synchronized HH neurons. (A) For the coupling coefficients *ε* = 1*e* − 4(i), 1*e* − 3(ii), 5*e* − 3(iii) and 1*e* − 2(iv), phase-stimulus plots show new phases *φ*) of the restoration of oscillation, using the color scale (on the right), when one of the coupled HH neurons is perturbed with different perturbation strengths (*dV* ) at the old phase points (*ϕ*) of the limit cycle. These plots also serve as isochron maps where voltage perturbations leading to identical new phase values can be identified based on colour. The set of perturbation strengths and old phases which together lead to phase indeterminate situations or annihilation of oscillation is shown as a bounded black patch on the map. This region serves as a portal to the null space or the basin of attraction of the stable focus. Further, the regions shown in grey are associated with perturbations which neither annihilate nor restore the original limit cycle. Rather, the system enters a new limit cycle with much smaller phase space span. This new limit cycle is however closely seated around the stable focus. Here, the values of the *dV* are ±[0, 2, 5, 10, 15, 20, 30, 40, 50, 60]*mV* and *ϕ* is discretized with a step of 0.05. The overlaid black patch of null space portal has been created with finely discretized values of *dV* and *ϕ* with step-sizes of 1*mV* and 0.001, respectively. (B) Variation in the area of null space portals in the regions of positive (i) and negative (ii) perturbations of the phase-stimulus plot across increasing coupling coefficient *ε*. SN denotes a single isolated HH neuron. The area of the null space portal is computed by adding together the spans of phase indeterminate *ϕ* over *dV* discretized by the step-size of 1mV. Variation in the total area of null space portal (iii) in the phase-stimulus plot across increasing *ε*.

## Discussion

The present study examines the effect of coupling on the null space of the HH model within the bistable regime with a stable limit cycle and a stable focus in the phase space. Stronger coupling between two HH models regularly generating APs in perfect in-phase synchronization leads to reduced null space. However, higher coupling strength between perfect anti-phase synchronized HH models significantly expands the null space.

A natural question arises on the biological significance of the passive diffusive coupling between the HH models considered here. Although chemical synapses do not fit in this description, electrical synapses via. gap junctions [44]have been widely recognized to describe bidirectional diffusive coupling between membrane potentials. Following additional detail, only somatic and axonal gap junctions should be discussed as dendritic gap junctions [45] bear many downstream complications disabling the instantaneous bidirectionality. Gap junctions are prevalent between pyramidal neurons and GABAergic interneurons during early development stages of the mammalian brain [46, 47]. However, they massively disappear in pyramidal populations towards adulthood and remain relatively abundant in the pyramidal neurons of the CA3 region of the adult hippocampus [48, 49]. However, the gap junctions remain quite abundant in a variety of interneuron populations in the adult brain, including the fast-spiking parvalbumin (PV) basket cells [ 51], regular interneurons [51] and the somatostatin-expressing (SSP)low-threshold interneurons [52]. Gap junctions are very essential for generating high-frequency synchronized rhythms [52–54]. Given the abundance of gap junctions amongst inhibitory neurons, as well as their critical role in rhythmic network activity, our findings highlight the vulnerability of neuronal spiking dynamics depending on the nature of synchronization between neural partners. Collapse into quiescent state due to perturbations can dramatically break the rhythm in local cluster of neurons. Therefore, strongly in-phase synchronized interneurons can avoid such collapse as they lack a null space in their phase space.

Another important point to note is that chemical synapses cannot be efficient in delivering sufficient voltage perturbations in soma and AIS for transitions between null space quiescent state and regular spiking in vivo. Chemical synaptic signals are slow time scale and undergo dendritic processing to ever cause large instantaneous voltage jumps. The perturbations must be strong and directly approachable to AP-generating segments and/or arising within the soma and AIS. Intrinsic stochasticity of the *Na*^+^ and *K*^+^ ion channels [55] can be one such factor locally within the soma/AIS. Moreover, spikelets transmitted through gap junctions [56, 57] and ephaptic coupling in tightly-packed neuronal surroundings [58] can be potential voltage perturbations as discussed in the present study. However, as external factors, brief optogenetic stimulation targeted to the soma or axon initial segment [59] can induce rapid membrane depolarizations that effectively act as voltage displacements on the timescale of spike initiation. Similarly, extracellular electrical stimulation [60] and local electric field perturbations [61] directly alter the somatic and axonal transmembrane potential.

A contrast between the hyperpolarized stable node discussed here and the other well-known phenomenon of depolarization block [62–64] needs to established. Both leads to a quiescent state where neurons are not able to spike despite strong background stimulation. The quiescent state under hyperpolarized stable node is long-lasting unless perturbation regenerates regular spiking by returning system onto the limit cycle. However, depolarization block is relatively transient with a gradual spontaneous return to regular spiking behaviour. This difference is further due to the underlying differences in the ionic mechanisms. Hyperpolarized stable node coexisting with a spiking limit cycle arises from a specific balance between inward and outward ionic currents in the spike-generating compartment of the neuron. In this regime, fast voltage-gated sodium channels are fully available but remain largely closed because the membrane potential is stabilized below spike threshold by dominant outward currents. Primarily delayed-rectifier potassium currents and, in many neurons, subthreshold potassium conductances such as leak K or low-threshold Kv channels contribute to this behaviour. Sodium inactivation is minimal, rendering the neuron excitable and capable of transitioning into repetitive spiking when sufficiently perturbed. In contrast, depolarization block emerges under sustained depolarizing drive, leading to substantial inactivation of sodium channels [62, 63]. Although outward potassium currents remain active, the loss of available Na channels prevents the system from sustaining a limit cycle, and the neuron settles into a depolarized, non-excitable equilibrium.

Electrical synapses via. gap junctions can be asymmetric as well [65, 66]. Moreover, they can undergo plasticity in the presence of neuromodulators such as dopamine as well as chemical signals such as nitric oxides [ 68]. These remain interesting aspects to be further explored in the context of null space associated with hyperpolarized quiescent state in the spiking dynamics. Our findings suggest a plausible role of gap junction plasticity and neuromodulation in tuning the vulnerability towards quiescent state. Incorporating this single neuron behaviour to neural network simulations would be highly beneficial in understanding how coupling strength and synchronization amongst inhibitory neurons with stochastic transitions between quiescent and spiking states can lead to dynamic rearrangements of network configuration and rhythm. Such a study can have strong implications in understanding neurodevelopmental and pathophysiological roles of rhythms in the presence of neuronal vulnerability to quiescent state.

## Supplementary Information

**Supplementary Table 1 and Supplementary Figures 1-12**

## Acknowledgments

RG and RKB acknowledge the DBT funding in the Centre for Computational Biology and Bioinformatics (BIC) 2025-26 project to support this research.

## Supplementary Table

**Table 1.**
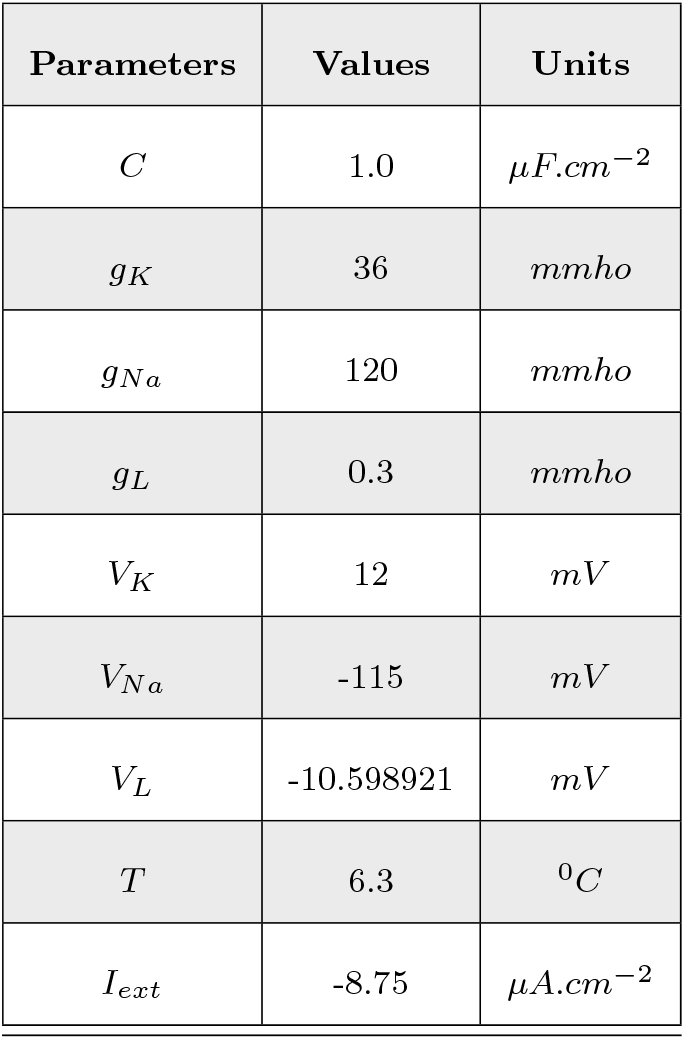
List of parameters present in the Hodgkin-Huxley model of a neuron. The values and units of the parameters used in the present study are provided here.

| Parameters | Values | Units |
| --- | --- | --- |
| $C$ | 1.0 | $\mu F.cm^{-2}$ |
| $g_K$ | 36 | $mmho$ |
| $g_{Na}$ | 120 | $mmho$ |
| $g_L$ | 0.3 | $mmho$ |
| $V_K$ | 12 | $mV$ |
| $V_{Na}$ | -115 | $mV$ |
| $V_L$ | -10.598921 | $mV$ |
| $T$ | 6.3 | $^{\circ}C$ |
| $I_{ext}$ | -8.75 | $\mu A.cm^{-2}$ |

## Supplementary Figures

**Figure 1.**
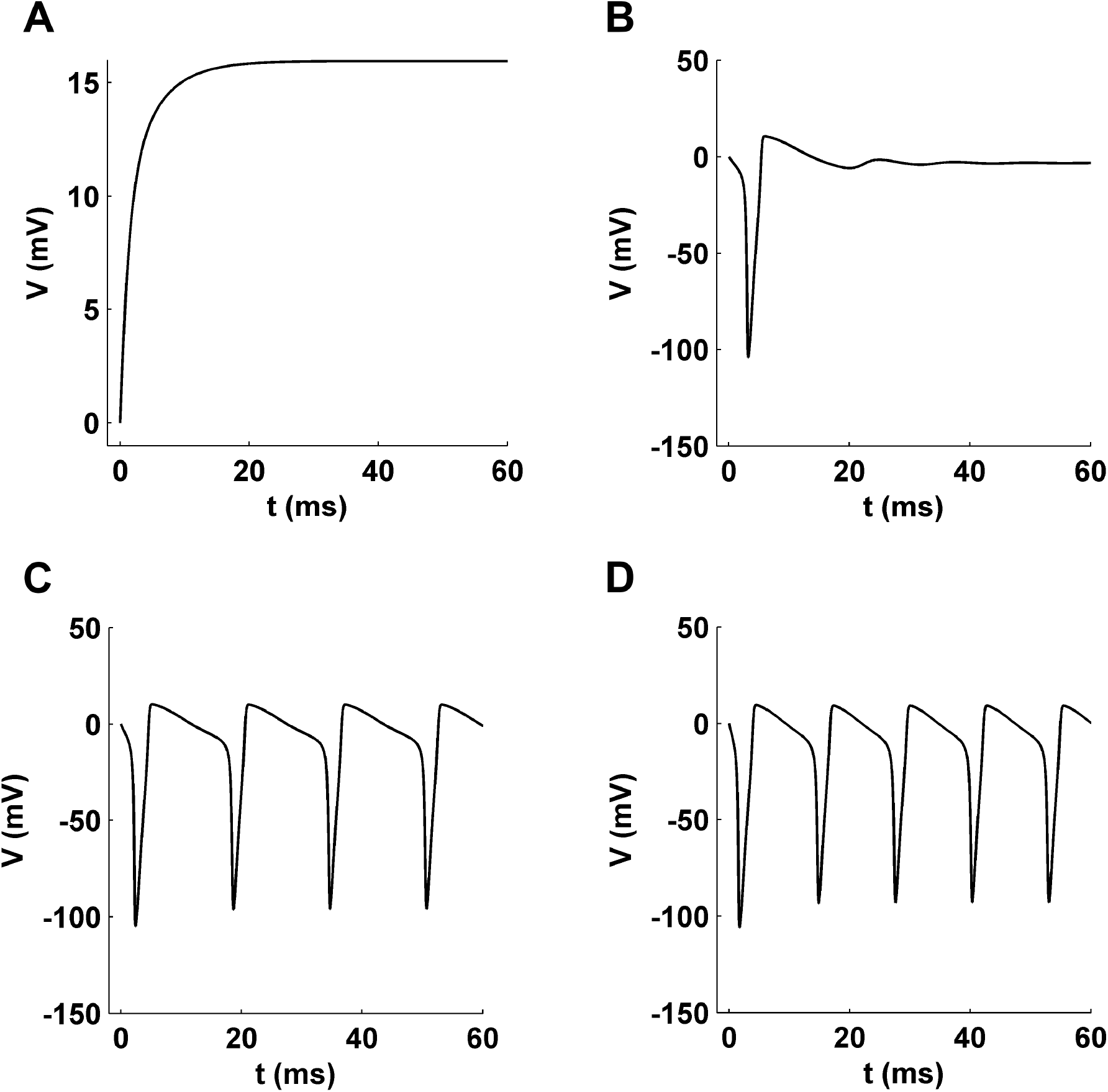
The responses of a HH neuron in terms of its membrane potential (V (t)) for different values of constant external current I_ext_. (A) For *I*_*ext*_ = 8.75*µA*.*cm*^−2^, the membrane potential exponentially rises to reach the stationary value of 15.9*mV* . (B) For *I*_*ext*_ = −5*µA*.*cm*^−2^, the membrane potential exhibits a damped oscillation and finally attains a stationary value of −4.5*mV* . The initial oscillation gives rise to a waveform of action potential. (C) For *I*_*ext*_ = −8.75*µA*.*cm*^−2^, the membrane potential regularly oscillates and results into repeated firing of action potentials by the HH neuron. (D) Similar repeated oscillation in the membrane potential is observed for stronger negative current, *I*_*ext*_ = −15*µA*.*cm*^−2^. The HH neuron is consistently studied at a temperature of 6.3^0^*C*.

**Figure 2.**
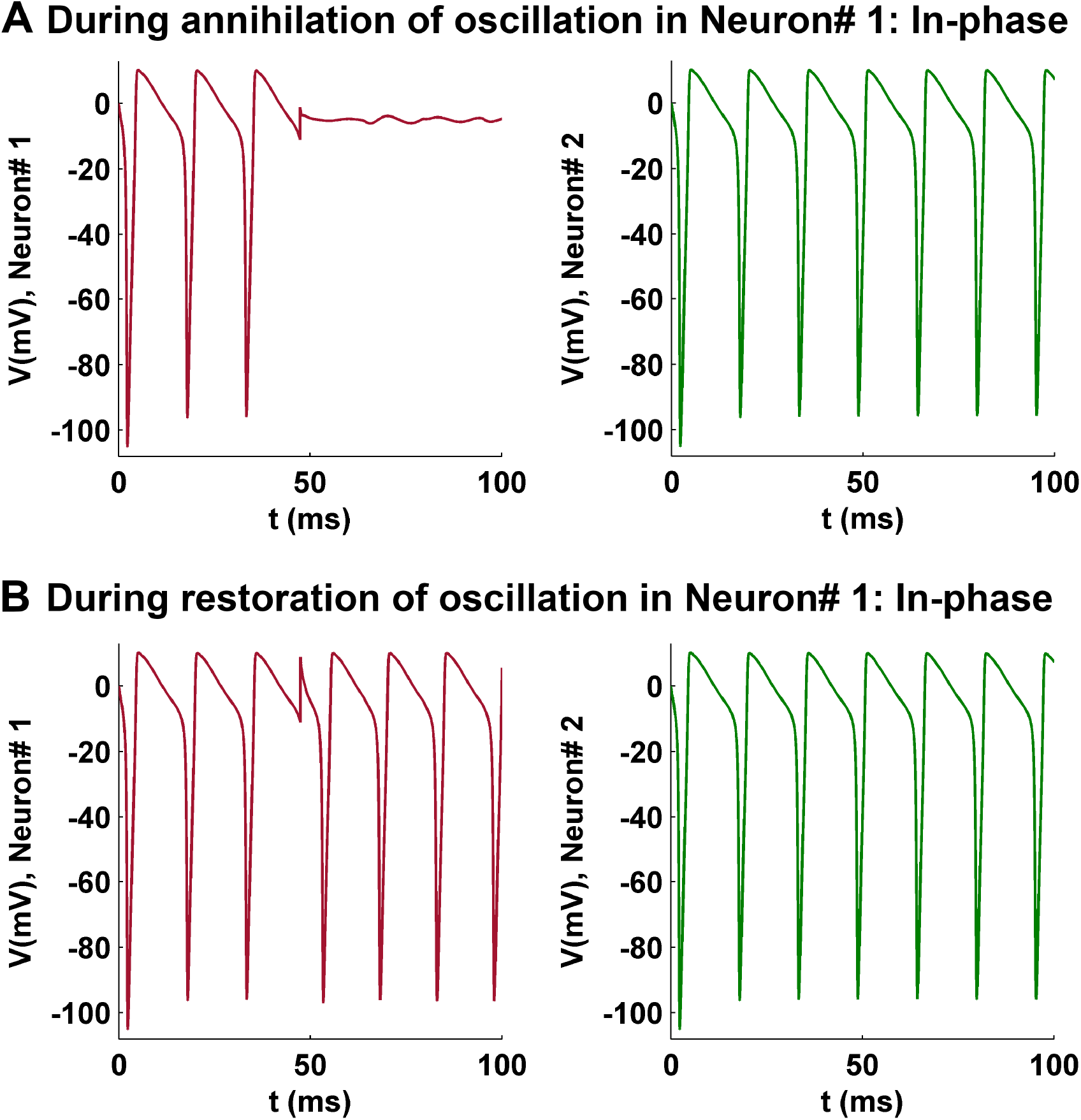
Effect of voltage perturbation on the oscillation of membrane potential in a coupled and in-phase synchronized system of two HH neurons. The two neurons are coupled with a sufficiently strong coupling coefficient, *ε* = 0.01. (A) Similar to a single HH neuron, perturbation of strength *dV* = 10*mV* at *ϕ* = 0.5 in Neuron #1 leads to the annihilation of the membrane potential oscillation. However, the oscillation in Neuron #2 remains completely unaffected from this event. (B) Perturbation of strength *dV* = 20*mV* at the same old phase *ϕ* = 0.5 leads to restoration of the oscillation in Neuron #1. However, Neuron #2 again hardly shows any change in the oscillation profile of its membrane potential.

**Figure 3.**
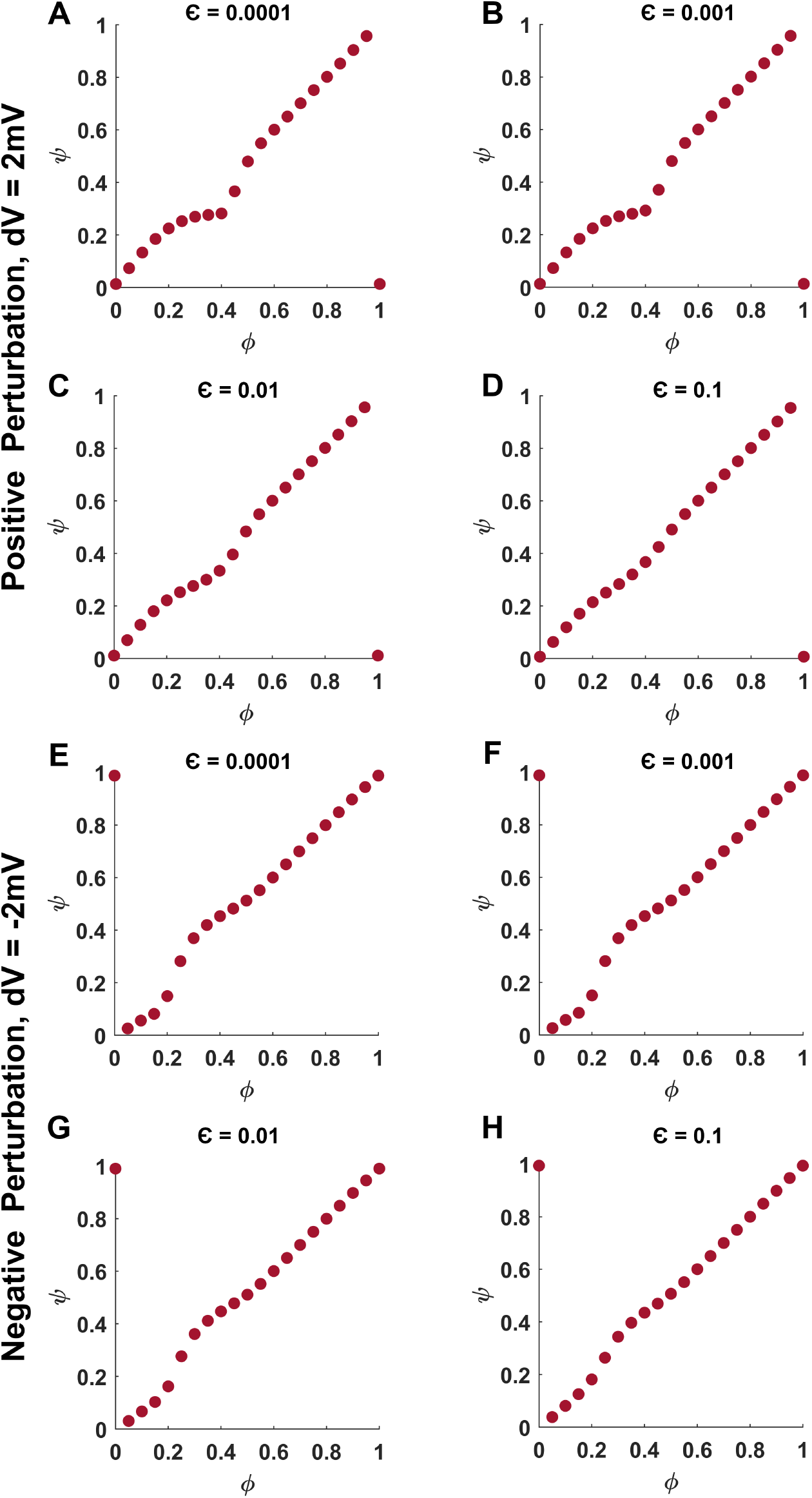
Phase response curve of a coupled and in-phase synchronized system of two HH neurons for absolute perturbation strength of 2mV. (A-D) The PRCs for the four coupling coefficients are shown in response to perturbation of strength *dV* = 2*mV* . (E-H) Similarly, the PRCs are shown in response to perturbation of strength *dV* = −2*mV* . All the PRCs belong to type 1 category.

**Figure 4.**
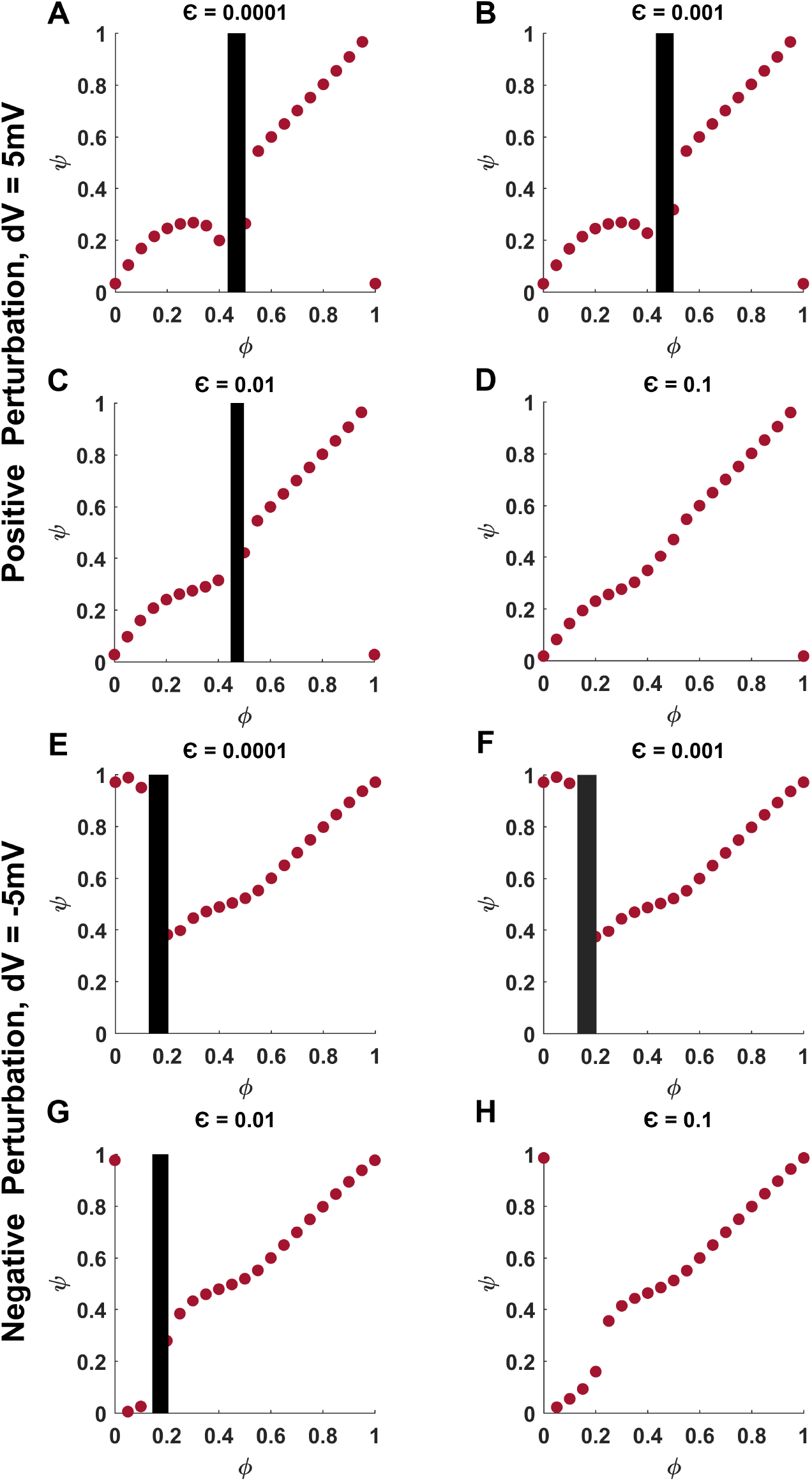
Phase response curve of a coupled and in-phase synchronized system of two HH neurons for absolute perturbation strength of 5mV. (A-D) The PRCs for the four coupling coefficients are shown in response to perturbation of strength *dV* = 5*mV* . (E-H) Similarly, the PRCs are shown in response to perturbation of strength *dV* = −5*mV* . All the PRCs belong to type 0 category, except (D) and (H) belonging to type 1 category.

**Figure 5.**
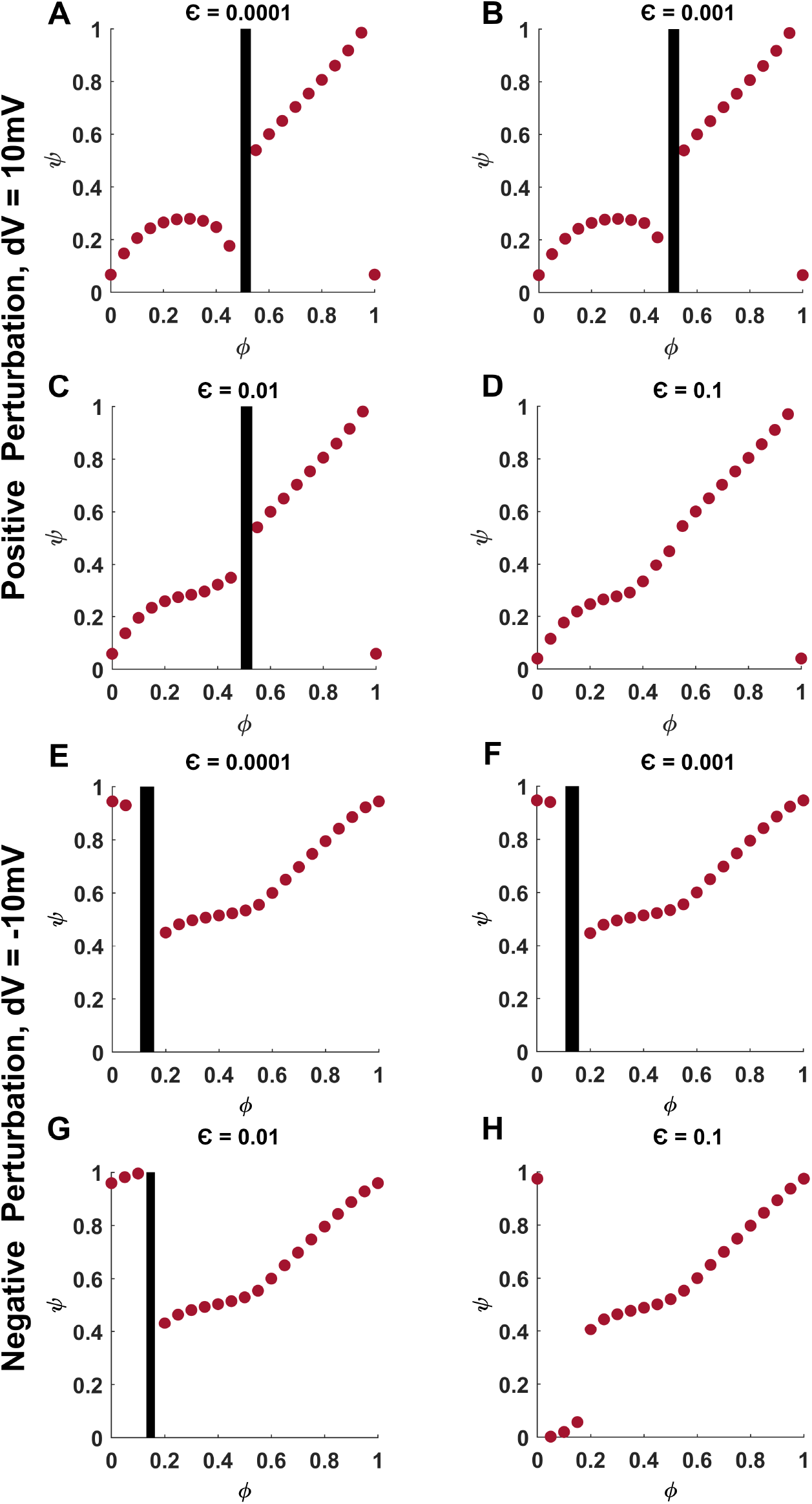
Phase response curve of a coupled and in-phase synchronized system of two HH neurons for absolute perturbation strength of 10mV. (A-D) The PRCs for the four coupling coefficients are shown in response to perturbation of strength *dV* = 10*mV* . (E-H) Similarly, the PRCs are shown in response to perturbation of strength *dV* = −10*mV* . All the PRCs belong to type 0 category, except (D) belonging to type 1 category.

**Figure 6.**
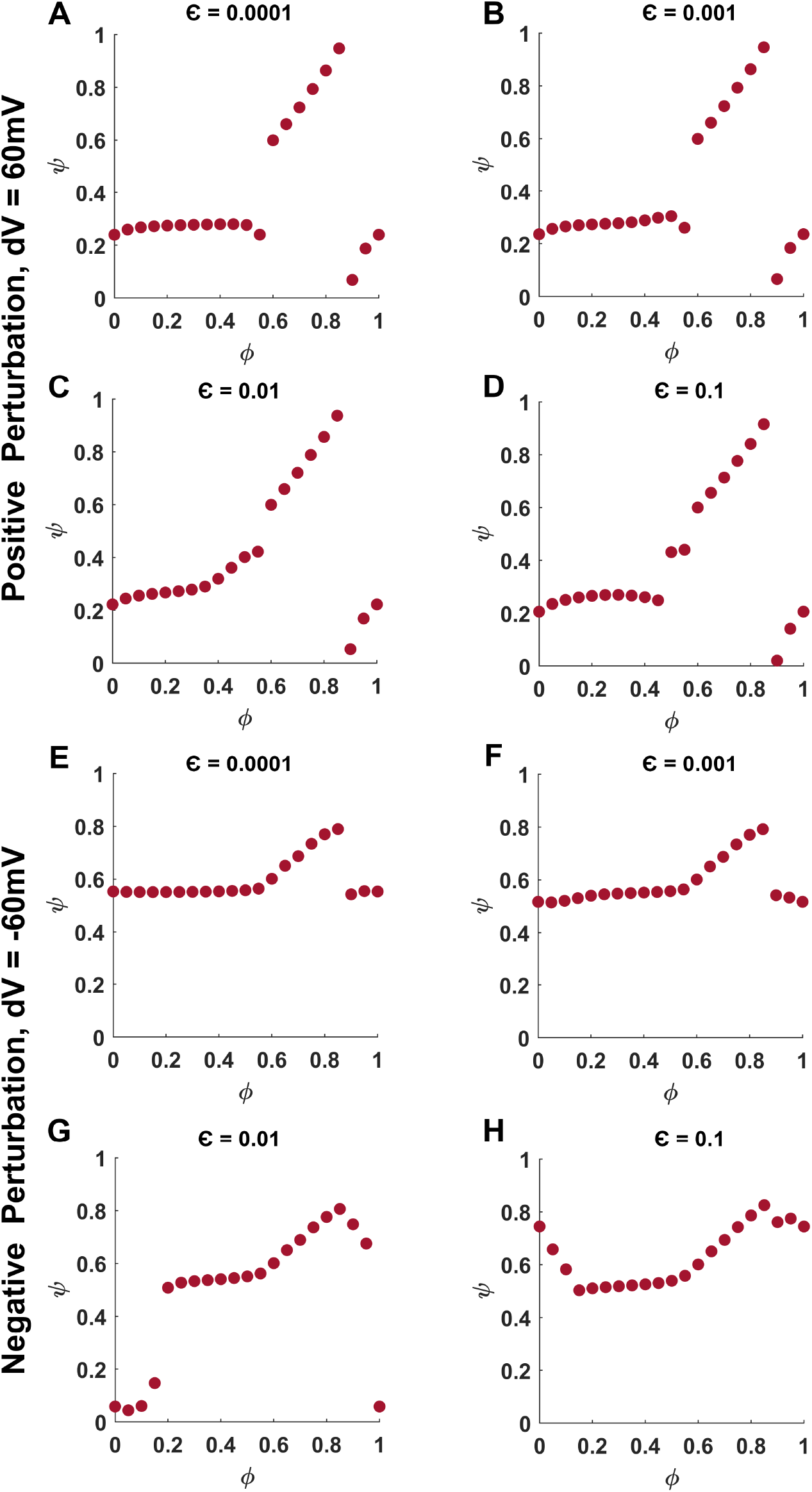
Phase response curve of a coupled and in-phase synchronized system of two HH neurons for absolute perturbation strength of 60mV. (A-D) The PRCs for the four coupling coefficients are shown in response to perturbation of strength *dV* = 60*mV* . (E-H) Similarly, the PRCs are shown in response to perturbation of strength *dV* = −60*mV* . All the PRCs belong to type 0 category.

**Figure 7.**
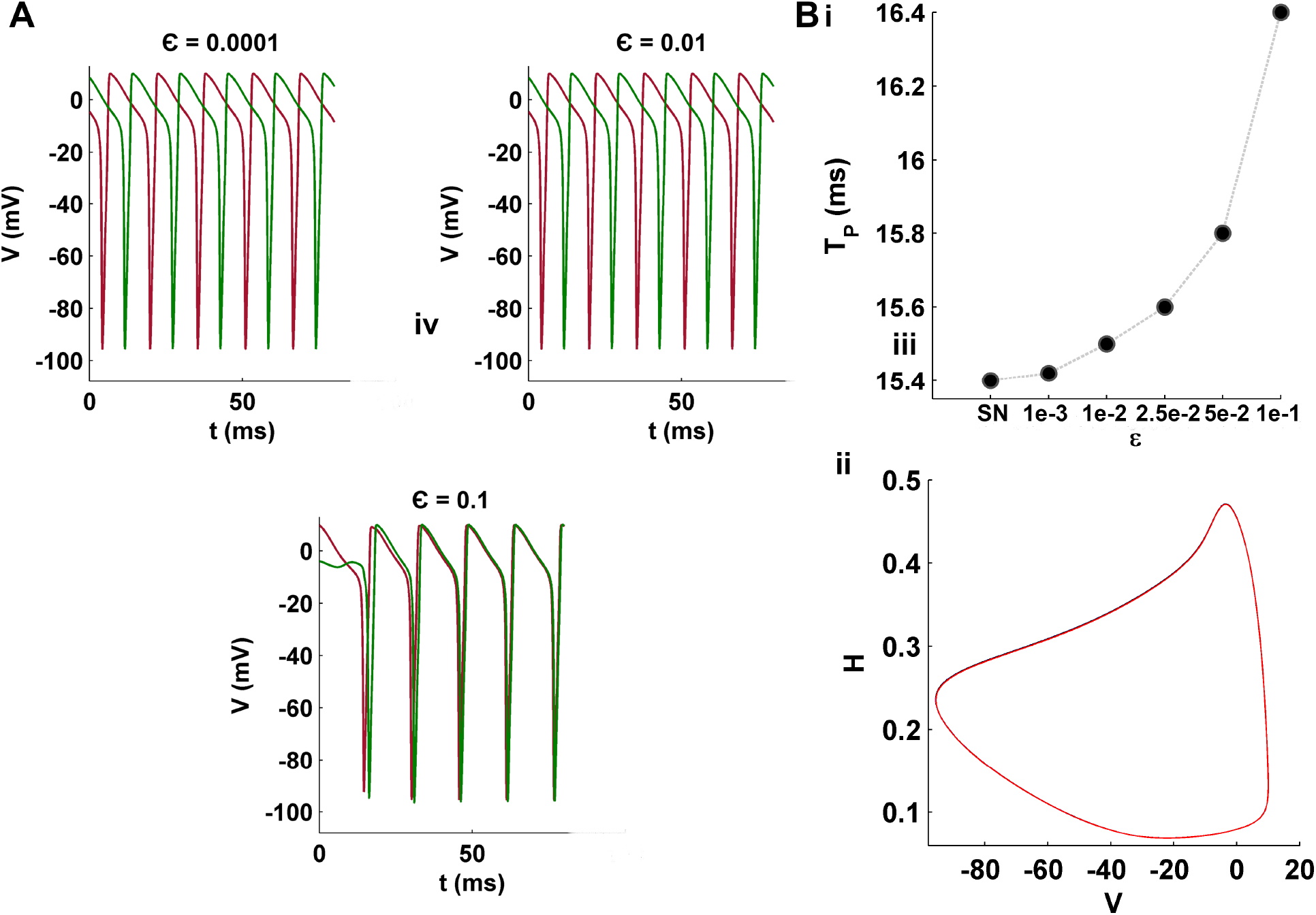
Effect of coupling coefficient on the stability of anti-phase synchronization and the time-period of membrane potential oscillation. (A) The perfect anti-phase synchronization between two diffusively-coupled HH neurons is stable only for the values of coupling coefficients *ε ≤* 1*e* − 2. For higher *ε*, the anti-phase synchronization spontaneously collapses into perfect in-phase synchronization. The perfect anti-phase synchronization is characterized with minimum cross-correlation of -0.2259 achievable between the temporal profiles of two HH neurons. (B) The time-period *T*_*P*_ of one cycle of oscillation in the membrane potential (i) gradually increases with the rise in coupling coefficient. SN represents the case of single HH neuron. The x-axis is not to scale. However, the limit cycles of oscillation (ii, overlaid in the *V* −*H* phase space) for different coupling coefficients remain intact and identical to that of single HH neuron.

**Figure 8.**
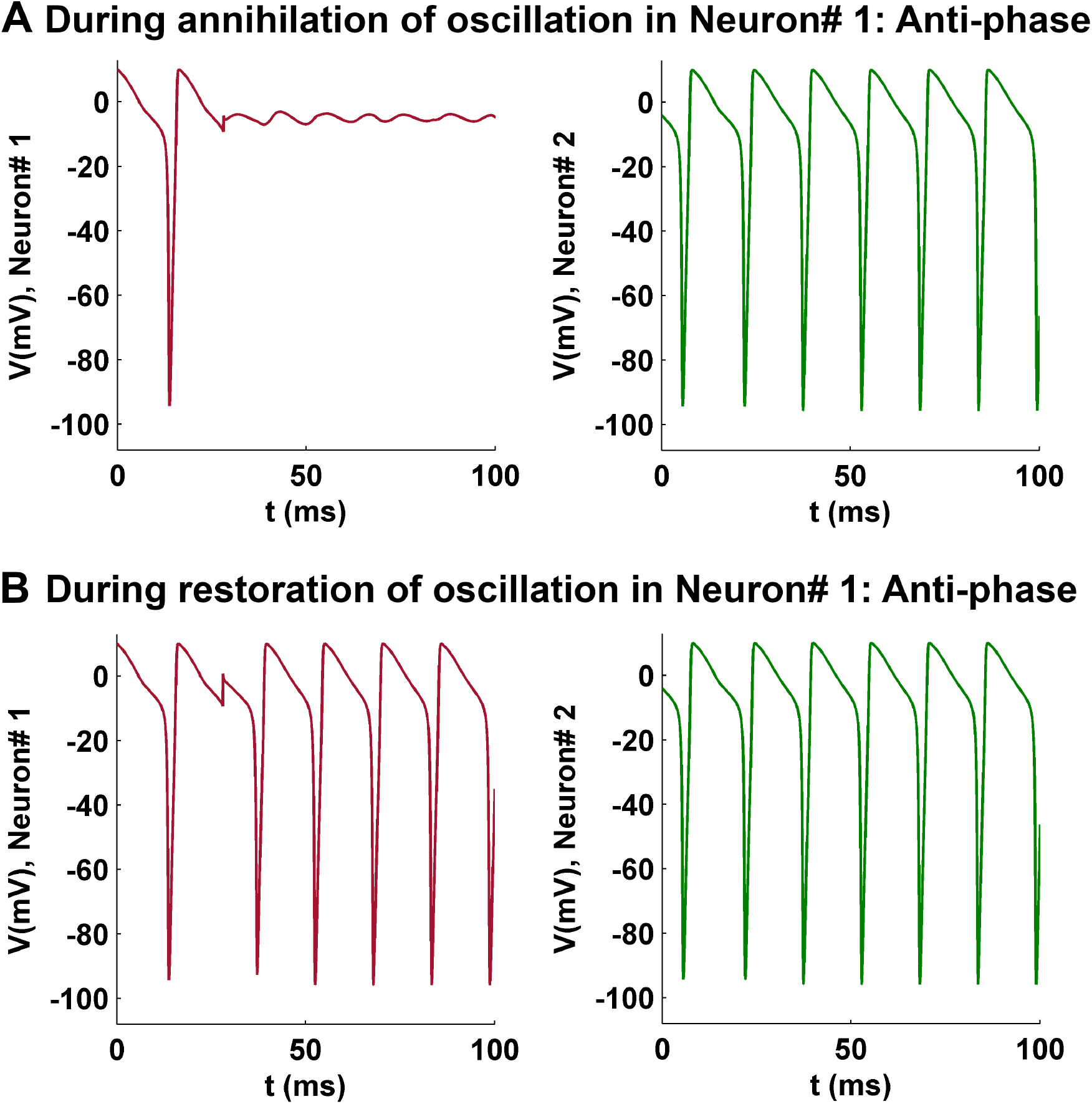
Effect of voltage perturbation on the oscillation of membrane potential in a coupled and anti-phase synchronized system of two HH neurons. The two neurons are coupled with a sufficiently strong coupling coefficient, *ε* = 0.01. (A) Perturbation of strength *dV* = 5*mV* at *ϕ* = 0.5 in Neuron #1 leads to the annihilation of the membrane potential oscillation. However, the oscillation in Neuron #2 remains completely unaffected from this event. (B) Perturbation of strength *dV* = 10*mV* at *ϕ* = 0.5 leads to restoration of the oscillation in Neuron #1. However, Neuron #2 again hardly shows any change in the oscillation profile of its membrane potential.

**Figure 9.**
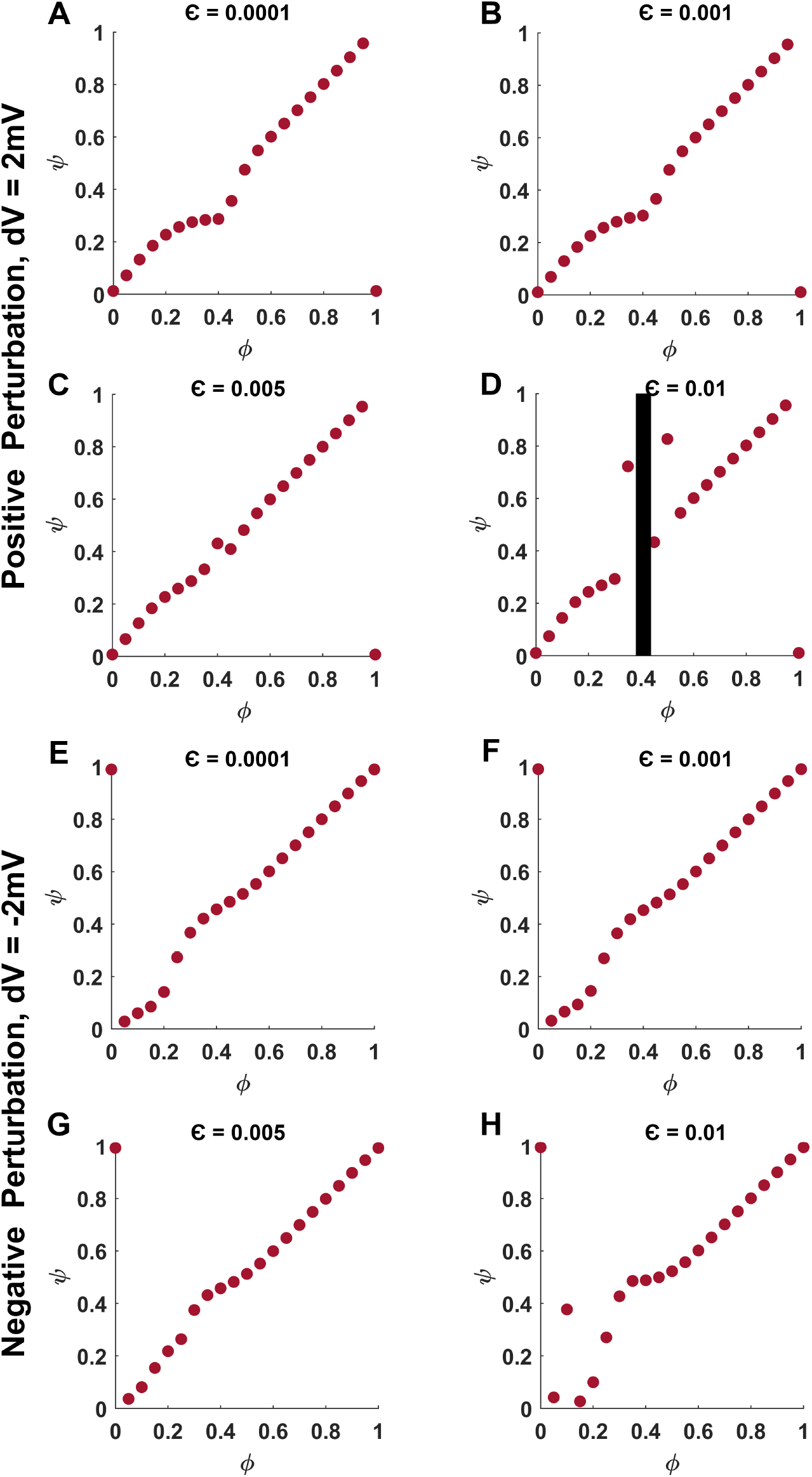
Phase response curve of a coupled and anti-phase synchronized system of two HH neurons for absolute perturbation strength of 2mV. (A-D) The PRCs for the four coupling coefficients are shown in response to perturbation of strength *dV* = 2*mV* . (E-H) Similarly, the PRCs are shown in response to perturbation of strength *dV* = −2*mV* . All the PRCs belong to type 1 category, except (C),(D), and (H) belonging to the type 0 category.

**Figure 10.**
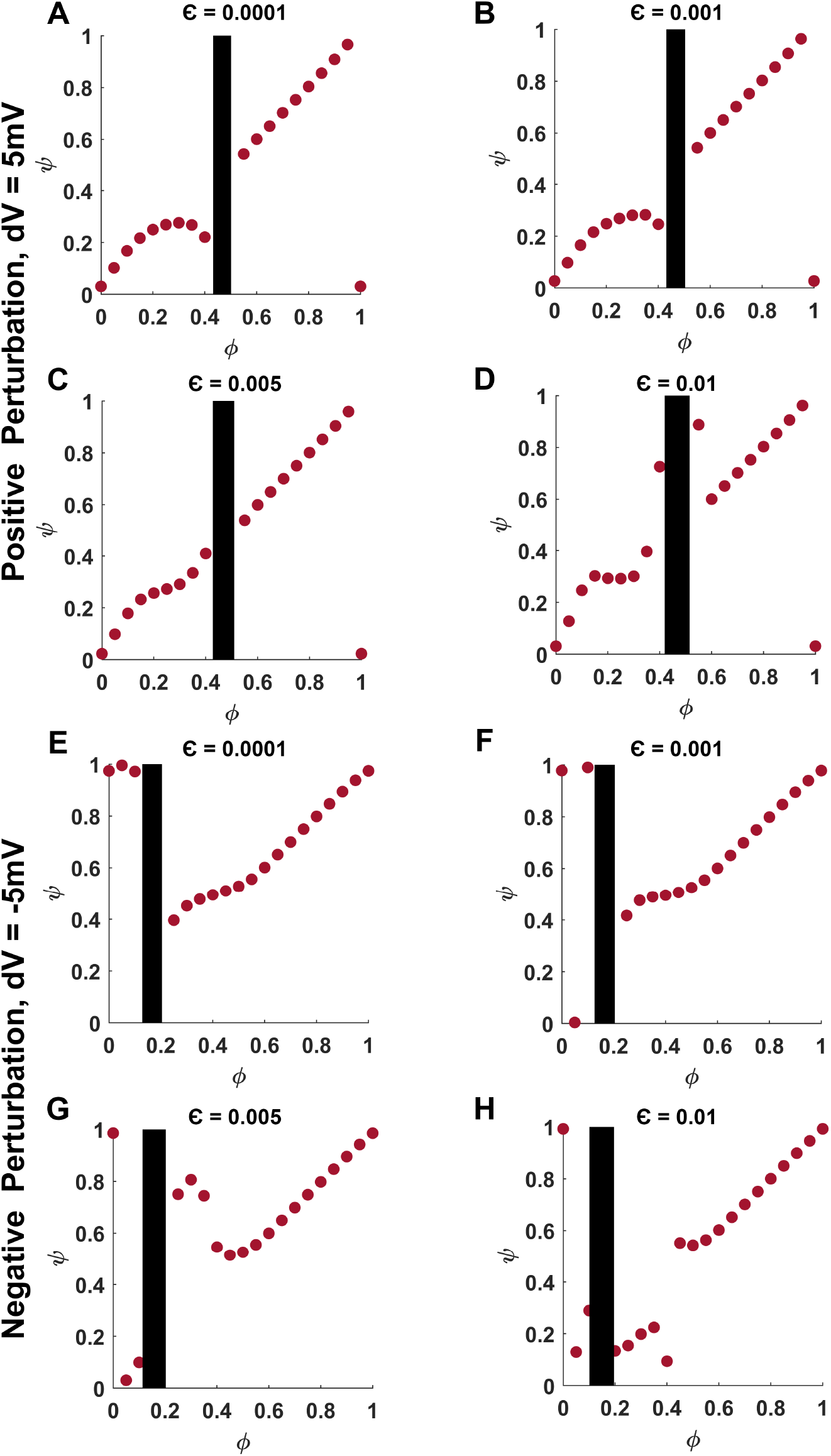
Phase response curve of a coupled and anti-phase synchronized system of two HH neurons for absolute perturbation strength of 5mV. (A-D) The PRCs for the four coupling coefficients are shown in response to perturbation of strength *dV* = 5*mV* . (E-H) Similarly, the PRCs are shown in response to perturbation of strength *dV* = −5*mV* . All the PRCs belong to type 0 category.

**Figure 11.**
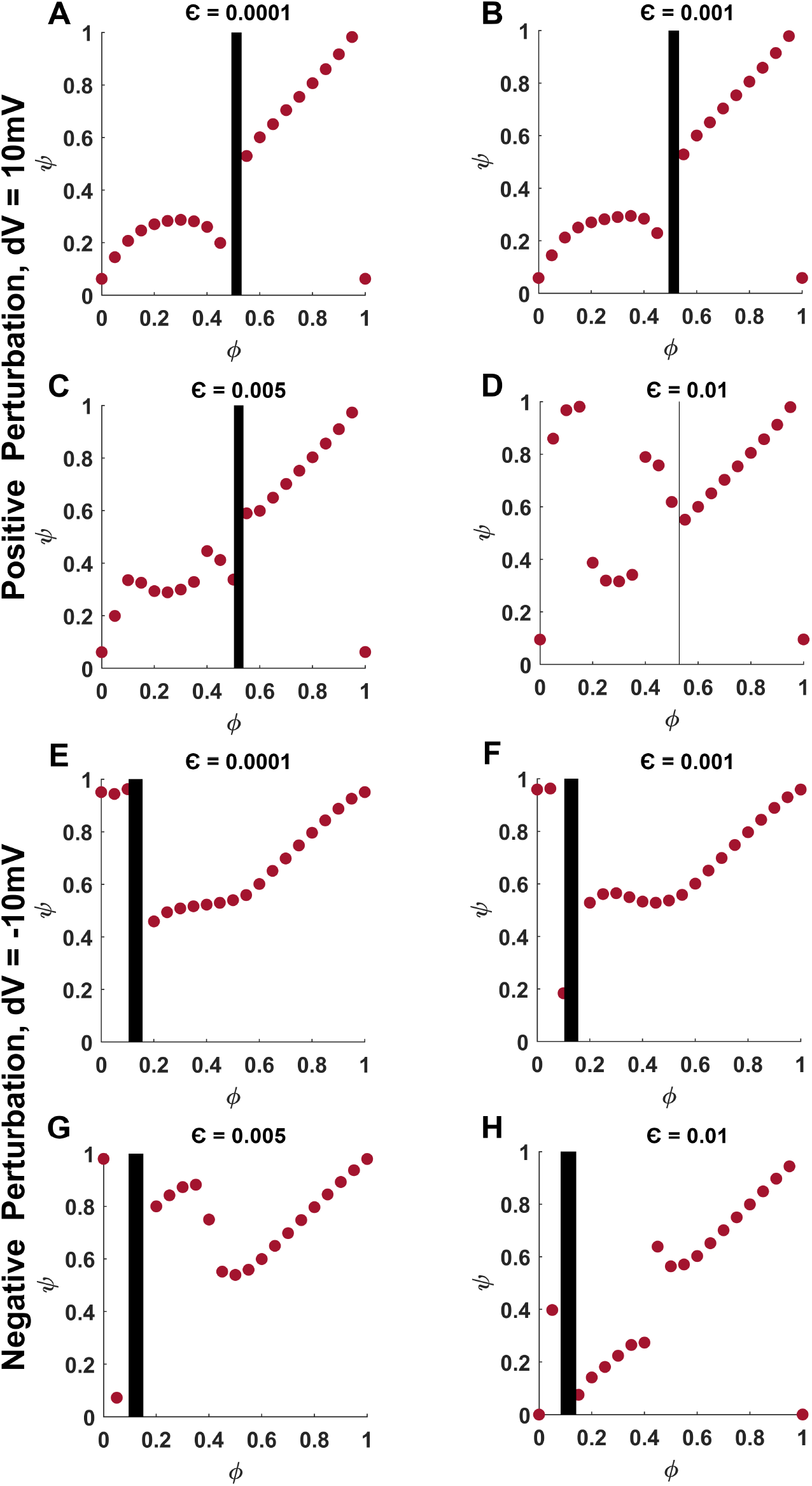
Phase response curve of a coupled and anti-phase synchronized system of two HH neurons for absolute perturbation strength of 10mV. (A-D) The PRCs for the four coupling coefficients are shown in response to perturbation of strength *dV* = 10*mV* . (E-H) Similarly, the PRCs are shown in response to perturbation of strength *dV* = −10*mV* . All the PRCs belong to type 0 category.

**Figure 12.**
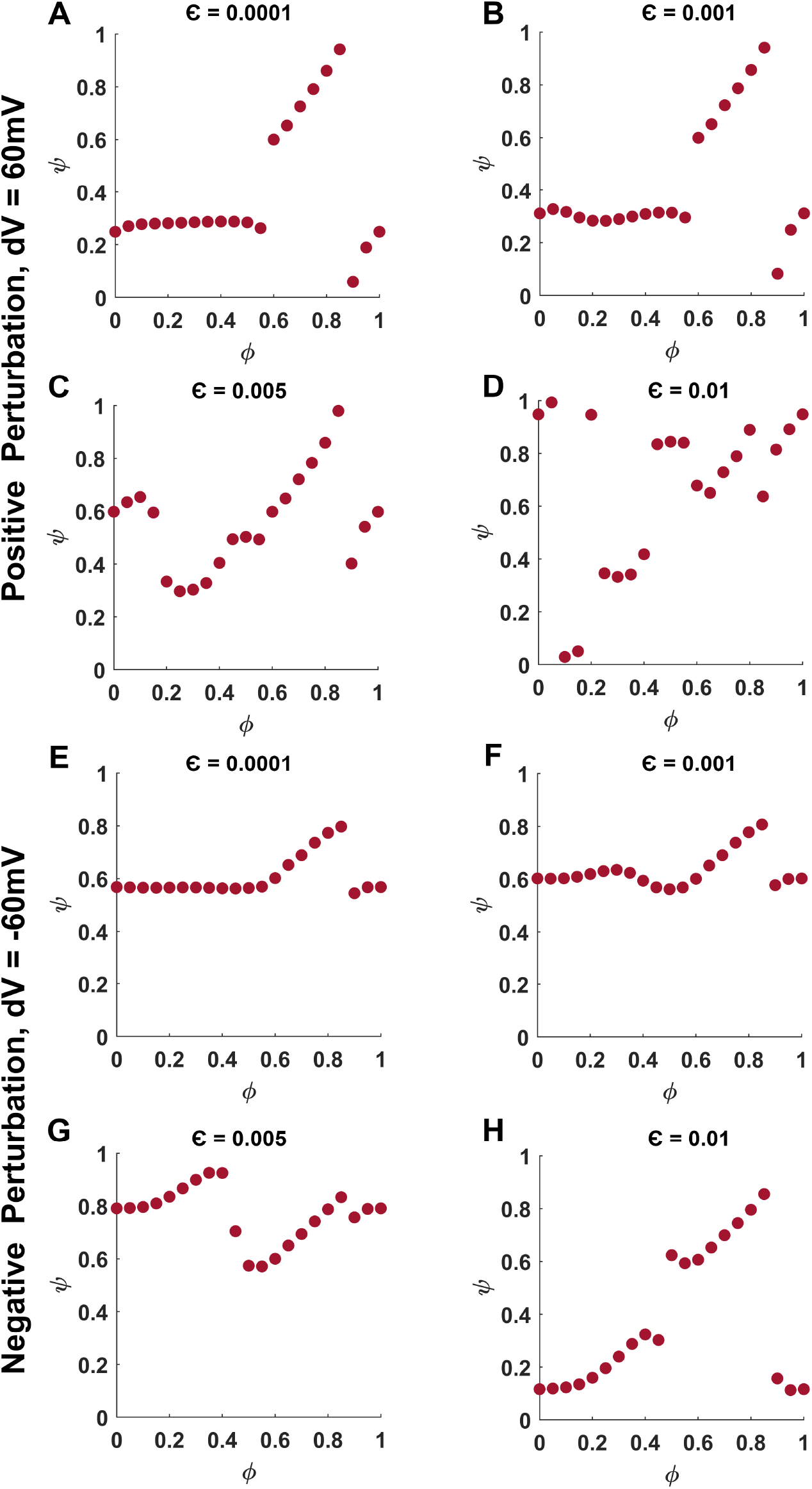
Phase response curve of a coupled and anti-phase synchronized system of two HH neurons for absolute perturbation strength of 60mV. (A-D) The PRCs for the four coupling coefficients are shown in response to perturbation of strength *dV* = 60*mV* . (E-H) Similarly, the PRCs are shown in response to perturbation of strength *dV* = −60*mV* . All the PRCs belong to type 0 category.

## References

1. Hodgkin, A.L., Huxley, A.F., Katz, B. (1952) Measurement of current voltage relations in the membrane of the giant axon of Loligo. Journal of Physiology 116, 242–448.

2. Hodgkin, A.L., Huxley, A.F. (1952) A quantitative description of membrane current and application to conduction and excitation in nerve. Journal of Physiology 117, 500–544.

3. Bean, B.P., 2007. The action potential in mammalian central neurons. Nature Reviews Neuroscience, 8(6), pp.451–465.

4. Izhikevich, E.M. (2007) Dynamical systems in neuroscience: the geometry of excitability and bursting. MIT Press, London, England.

5. Gerstner, W., Kistler, W.M., Naud, R. and Paninski, L., 2014. Neuronal dynamics: From single neurons to networks and models of cognition. Cambridge University Press.

6. Petousakis, K.E., Apostolopoulou, A.A. and Poirazi, P., 2023. The impact of HodgkinHuxley models on dendritic research. Journal of physiology, 601(15), pp.3091–3102.

7. Kole, M.H. and Stuart, G.J., 2012. Signal processing in the axon initial segment. Neuron, 73(2), pp.235–247.

8. Hfflin, F., Jack, A., Riedel, C., Mack-Bucher, J., Roos, J., Corcelli, C., Schultz, C., Wahle, P. and Engelhardt, M., 2017. Heterogeneity of the axon initial segment in interneurons and pyramidal cells of rodent visual cortex. Frontiers in cellular neuroscience, 11, p.332.

9. Kamaleddin, M.A., 2025. Biophysical properties of the membrane influence spike initiation dynamics and neuronal excitability: a focus on Kv1 channels in myelinated axons. Proceedings of the Royal Society B: Biological Sciences, 292(2051).

10. Guttman, R., Barnhill, R. (1970) Oscillation and repetitive firing in squid axons. Journal of General Physiology, 55, 104–118.

11. Guttman, R., Lewis, S. (1978) Repetitive firing in squid axon membrane as a model for a neuronal oscillator. Biology Bulletin, 155, 441.

12. Guttman, R., Lewis, S., Rinzel, J. (1979) Control of repetitive firing in squid axon membrane as a model for a neuron oscillator. Journal of Physiology, 305, 377–395.

13. Aihara, K. and Matsumoto, G., 1983. Two stable steady states in the Hodgkin-Huxley axons. Biophysical journal, 41(1), pp.87–89.

14. Luccioli, S., Kreuz, T. and Torcini, A., 2006. Dynamical response of the Hodgkin-Huxley model in the high-input regime. Physical Review EStatistical, Nonlinear, and Soft Matter Physics, 73(4), p.041902.

15. Rowat, P.F. and Greenwood, P.E., 2014. The ISI distribution of the stochastic Hodgkin-Huxley neuron. Frontiers in Computational Neuroscience, 8, p.111.

16. Kameneva, T., Meffin, H., Burkitt, A.N. and Grayden, D.B., 2018, July. Bistability in Hodgkin-Huxley-type equations. In 2018 40th Annual International Conference of the IEEE Engineering in Medicine and Biology Society (EMBC) (pp. 4728–4731). IEEE.

17. Lu, Y., Xin, X. and Rinzel, J., 2023. Bistability at the onset of neuronal oscillations. Biological cybernetics, 117(1), pp.61–79.

18. Williams, S.R., Christensen, S.R., Stuart, G.J. and Husser, M., 2002. Membrane potential bistability is controlled by the hyperpolarization-activated current IH in rat cerebellar Purkinje neurons in vitro. The Journal of physiology, 539(2), pp.469–483.

19. Loewenstein, Y., Mahon, S., Chadderton, P., Kitamura, K., Sompolinsky, H., Yarom, Y. and Husser, M., 2005. Bistability of cerebellar Purkinje cells modulated by sensory stimulation. Nature neuroscience, 8(2), pp.202–211.

20. Plotkin, J.L., Day, M. and Surmeier, D.J., 2011. Synaptically driven state transitions in distal dendrites of striatal spiny neurons. Nature neuroscience, 14(7), pp.881–888.

21. Rekling, J.C., Funk, G.D., Bayliss, D.A., Dong, X.W. and Feldman, J.L., 2000. Synaptic control of motoneuronal excitability. Physiological reviews, 80(2), pp.767–852.

22. Wiegert, J.S., Mahn, M., Prigge, M., Printz, Y. and Yizhar, O., 2017. Silencing neurons: tools, applications, and experimental constraints. Neuron, 95(3), pp.504–529.

23. Malashchenko, T., Shilnikov, A. and Cymbalyuk, G., 2011. Bistability of bursting and silence regimes in a model of a leech heart interneuron. Physical Review EStatistical, Nonlinear, and Soft Matter Physics, 84(4), p.041910.

24. Winfree, A.T. (1973) Time and timelessness in biological clocks. In temporal aspects of therapeutics. Vol. 2, Plenum Publishing Corporation, New York.

25. Winfree, A.T. (1974) Patterns of phase compromise in biological cycles. Journal of Mathematical Biology, 1, 73–95.

26. Winfree, A.T. (1972) Oscillatory glycolysis in yeast: the pattern of phase resetting oxygen. Archives of Biochemistry and Biophysics, 149, 388–401.

27. Winfree, A.T. (1971) Corkscrews and singularities in fruitflies: resetting behavior of the circadian eclosion rhythm. Biochronometry. National Academy of Sciences, Washington, D.C. 81–109.

28. Guckenheimer, J. (1975) Isochrons and phaseless sets. Journal of Mathematical Biology, 1, 259–273.

29. Best, E.N. (1979) Null space in the Hodgkin-Huxley equations. A critical test. Biophysical Journal, 27, 87–104.

30. Pikvosky, A., Rosenblum, M., Kurths, J. (2001) Synchronization. A universal concept in nonlinear sciences. Cambridge University Press, UK.

31. Elson, R.C., Selverston, A.I., Huerta, R., Rulkov, N.F., Rabinovich, M.I., Abarbanel, H.D.I. (1998) Synchronous behavior of two coupled biological neurons. Physics Review Letters, 81, 5692–5695.

32. Pinto, R.D., Varona, P., Volkovskii, R., Abarbanel, H.D.I., Rabinovich, M.I. (2000) Synchronous behavior of two coupled electronic neurons. Physical Review E, 62, 2644–2656.

33. Smeal, R.M., Ermentrout, G.B. and White, J.A., 2010. Phase-response curves and synchronized neural networks. Philosophical Transactions of the Royal Society B: Biological Sciences, 365(1551), pp.2407–2422.

34. Marwan, N. and Kurths, J., 2002. Nonlinear analysis of bivariate data with cross recurrence plots. Physics Letters A, 302(5-6), pp.299–307.

35. Shockley, K., Butwill, M., Zbilut, J.P. and Webber Jr, C.L., 2002. Cross recurrence quantification of coupled oscillators. Physics Letters A, 305(1-2), pp.59–69.

36. Romano, M.C., Thiel, M., Kurths, J. and von Bloh, W., 2004. Multivariate recurrence plots. Physics letters A, 330(3-4), pp.214–223.

37. Marwan, N., Romano, M.C., Thiel, M. and Kurths, J., 2007. Recurrence plots for the analysis of complex systems. Physics reports, 438(5-6), pp.237–329.

38. Goswami, B., 2019. A brief introduction to nonlinear time series analysis and recurrence plots. Vibration, 2(4), pp.332–368.

39. Webber Jr, C.L. and Zbilut, J.P., 1994. Dynamical assessment of physiological systems and states using recurrence plot strategies. Journal of applied physiology, 76(2), pp.965–973.

40. Zbilut, J.P., Giuliani, A. and Webber Jr, C.L., 1998. Detecting deterministic signals in exceptionally noisy environments using cross-recurrence quantification. Physics Letters A, 246(1-2), pp.122–128.

41. Babaei, B., Zarghami, R., Sedighikamal, H., Sotudeh-Gharebagh, R. and Mostoufi, N., 2014. Selection of minimal length of line in recurrence quantification analysis. Physica A: Statistical Mechanics and its Applications, 395, pp.112–120.

42. Ao, X., Hänggi, P., Schmid, G. (2013) In-phase and anti-phase synchronization in noisy Hodgkin-Huxley neurons. Mathematical Biosciences, 245, 49–55.

43. Sterratt, D.C., 2022. Q10: the effect of temperature on ion channel kinetics. In Encyclopedia of Computational Neuroscience (pp. 2949–2950). New York, NY: Springer New York.

44. Shl, G., Maxeiner, S. and Willecke, K., 2005. Expression and functions of neuronal gap junctions. Nature reviews neuroscience, 6(3), pp.191–200.

45. Traub, R.D., Pais, I., Bibbig, A., LeBeau, F.E., Buhl, E.H., Hormuzdi, S.G., Monyer, H. and Whittington, M.A., 2003. Contrasting roles of axonal (pyramidal cell) and dendritic (interneuron) electrical coupling in the generation of neuronal network oscillations. Proceedings of the National Academy of Sciences, 100(3), pp.1370–1374.

46. Yuste, R., Peinado, A. and Katz, L.C., 1992. Neuronal domains in developing neocortex. Science, 257(5070), pp.665–669.

47. Sutor, B. and Hagerty, T., 2005. Involvement of gap junctions in the development of the neocortex. Biochimica et Biophysica Acta (BBA)-Biomembranes, 1719(1-2), pp.59–68.

48. Schmitz, D., Schuchmann, S., Fisahn, A., Draguhn, A., Buhl, E.H., Petrasch-Parwez, E., Dermietzel, R., Heinemann, U. and Traub, R.D., 2001. Axo-axonal coupling: a novel mechanism for ultrafast neuronal communication. Neuron, 31(5), pp.831–840.

49. Hamzei-Sichani, F., Kamasawa, N., Janssen, W.G., Yasumura, T., Davidson, K.G., Hof, P.R., Wearne, S.L., Stewart, M.G., Young, S.R., Whittington, M.A. and Rash, J.E., 2007. Gap junctions on hippocampal mossy fiber axons demonstrated by thin-section electron microscopy and freezefracture replica immunogold labeling. Proceedings of the National Academy of Sciences, 104(30), pp.12548–12553.

50. Galarreta, M. and Hestrin, S., 2002. Electrical and chemical synapses among parvalbumin fast-spiking GABAergic interneurons in adult mouse neocortex. Proceedings of the National Academy of Sciences, 99(19), pp.12438–12443.

51. Hatch, R.J., Mendis, G.D.C., Kaila, K., Reid, C.A. and Petrou, S., 2017. Gap junctions link regular-spiking and fast-spiking interneurons in layer 5 somatosensory cortex. Frontiers in Cellular Neuroscience, 11, p.204.

52. Gibson, J.R., Beierlein, M. and Connors, B.W., 1999. Two networks of electrically coupled inhibitory neurons in neocortex. Nature, 402(6757), pp.75–79.

53. Hjorth, J., Blackwell, K.T. and Kotaleski, J.H., 2009. Gap junctions between striatal fast-spiking interneurons regulate spiking activity and synchronization as a function of cortical activity. Journal of Neuroscience, 29(16), pp.5276–5286.

54. Pernelle, G., Nicola, W. and Clopath, C., 2018. Gap junction plasticity as a mechanism to regulate network-wide oscillations. PLoS computational biology, 14(3), p.e1006025.

55. Faisal, A.A., Selen, L.P.J., Wolpert, D.M. (2008) Noise in the nervous system. Nature Reviews Neuroscience, 9, 292–303.

56. Apostolides, P.F. and Trussell, L.O., 2013. Regulation of interneuron excitability by gap junction coupling with principal cells. Nature Neuroscience, 16(12), pp.1764–1772.

57. Michalikova, M., Remme, M.W., Schmitz, D., Schreiber, S. and Kempter, R., 2019. Spikelets in pyramidal neurons: generating mechanisms, distinguishing properties, and functional implications. Reviews in the Neurosciences, 31(1), pp.101–119.

58. Anastassiou, C.A., Perin, R., Markram, H. and Koch, C., 2011. Ephaptic coupling of cortical neurons. Nature neuroscience, 14(2), pp.217–223.

59. Grubb, M.S. and Burrone, J., 2010. Channelrhodopsin-2 localised to the axon initial segment. PloS one, 5(10), p.e13761.

60. Radivojevic, M., Jckel, D., Altermatt, M., Mller, J., Viswam, V., Hierlemann, A. and Bakkum, D.J., 2016. Electrical identification and selective microstimulation of neuronal compartments based on features of extracellular action potentials. Scientific reports, 6(1), p.31332.

61. Liu, A., Vrslakos, M., Kronberg, G., Henin, S., Krause, M.R., Huang, Y., Opitz, A., Mehta, A., Pack, C.C., Krekelberg, B. and Bernyi, A., 2018. Immediate neurophysiological effects of transcranial electrical stimulation. Nature communications, 9(1), p.5092.

62. Bianchi, D., Marasco, A., Limongiello, A., Marchetti, C., Marie, H., Tirozzi, B. and Migliore, M., 2012. On the mechanisms underlying the depolarization block in the spiking dynamics of CA1 pyramidal neurons. Journal of computational neuroscience, 33(2), pp.207–225.

63. Qian, K., Yu, N., Tucker, K.R., Levitan, E.S. and Canavier, C.C., 2014. Mathematical analysis of depolarization block mediated by slow inactivation of fast sodium channels in midbrain dopamine neurons. Journal of neurophysiology, 112(11), pp.2779–2790.

64. Calin, A., Ilie, A.S. and Akerman, C.J., 2021. Disrupting epileptiform activity by preventing parvalbumin interneuron depolarization block. Journal of Neuroscience, 41(45), pp.9452–9465.

65. Mendoza, A. and Haas, J.S., 2021. Electrical synapse asymmetry results from, and masks, neuronal heterogeneity. bioRxiv, pp.2021–06.

66. Vaughn, M.J. and Haas, J.S., 2022. On the diverse functions of electrical synapses. Frontiers in Cellular Neuroscience, 16, p.910015.

67. Haas, J.S., Greenwald, C.M. and Pereda, A.E., 2016. Activity-dependent plasticity of electrical synapses: increasing evidence for its presence and functional roles in the mammalian brain. BMC cell biology, 17(Suppl 1), p.14.

68. Coulon, P. and Landisman, C.E., 2017. The potential role of gap junctional plasticity in the regulation of state. Neuron, 93(6), pp.1275–1295.

